# The roles of GpsB and DivIVA in *Staphylococcus aureus* growth and division

**DOI:** 10.1101/2023.06.16.545239

**Authors:** Joshua A. F. Sutton, Mark Cooke, Mariana Tinajero-Trejo, Katarzyna Wacnik, Bartłomiej Salamaga, Callum Portman-Ross, Victoria A. Lund, Jamie K. Hobbs, Simon J. Foster

## Abstract

The spheroid bacterium *S. aureus* is often used as a model of morphogenesis due to its apparent simple cell cycle. *S. aureus* has many cell division proteins that are conserved across bacteria alluding to common functions. However, despite intensive study we still do not know the roles of many of these components. Here we have examined the functions of the paralogues DivIVA and GpsB in the *S. aureus* cell cycle. Cells lacking *gpsB* display a more spherical phenotype than wild type, associated with a decrease in peripheral cell wall peptidoglycan synthesis. This correlates with an increased localisation of penicillin binding proteins at the developing septum, notably PBPs 2 and 3. Our results highlight the role of GpsB as an apparent regulator of cell morphogenesis in *S*. *aureus*.

## Introduction

The cell envelope for most bacteria maintains cell shape and viability as well as forming an interface with the environment (Turner *et al*., 2014). In Gram-positive organisms the envelope consists of a cell membrane containing lipoteichoic acids (LTA) surrounded by thick layer of peptidoglycan (PG) decorated with wall teichoic acids (WTA) and surface proteins (Vollmer *et al*., 2008). The cell wall is dynamic, having to retain cellular integrity in the face of internal turgor while still permitting growth and division. In rod-shaped cells, the machinery required for vegetative growth is called the elongasome, while that for cell division is the divisome (Cabeen and Jacobs-Wagner, 2005). Even the spheroid bacterium *Staphylococcus aureus* shows some elongation of the cell during growth, but lacks an elongasome (Reichmann *et al*., 2019). Bacterial cell growth and division are highly organised and complex processes. PG structural dynamics are required for morphogenesis, with synthesis and hydrolysis being tightly controlled (Wheeler *et al*., 2015; Zhou *et al*., 2015). The final stages of PG synthesis are performed largely through the transglycosylase (TG) and transpeptidase (TP) activities of penicillin binding proteins (PBPs) (Typas *et al*., 2011). *S. aureus* encodes four native PBPs, where PBP1 (monofunctional TP) and PBP2 (bifunctional TG and TP) are essential (Pinho and Errington, 2005; Wacnik *et al*., 2022). PBP1 has multiple functions during cell division, both enzymatically and as a scaffold (Wacnik *et al*., 2022). PBP3 is a nonessential TP (Pinho *et al*., 2000). The monofunctional TGs FtsW and RodA, both shape, elongation, division, and sporulation (SEDS) proteins, form cognate pairs with PBP1 (responsible for septum formation) and PBP3 (responsible for peripheral PG synthesis and cell shape maintenance) respectively (Reichmann *et al*., 2019). PBP4 has D,D-carboxypeptidase and TP activity which results in the high level of PG crosslinking associated with *S. aureus* (Wyke *et al*., 1981; Atilano *et al*., 2010; Loskill *et al*., 2014). Methicillin resistant *Staphylococcus aureus* (MRSA) strains possess the additional TP PBP2a which has low-affinity for β-lactam antibiotics (Hartman and Tomasz, 1984; Pinho *et al*., 2001a).

Cell division is both spatially and temporally regulated to ensure the maintenance of cell shape and integrity. Staphylococcal cell division begins with the formation of the Z-ring, where multiple FtsZ monomers polymerise to form a scaffold to recruit divisome proteins that allow septation (Pinho *et al*., 2013; Szwedziak *et al*., 2014). Cell division must occur after DNA replication and subsequent chromosome segregation to ensure that septa do not split the separating chromosomes. In part this is achieved by proteins such as Noc (nucleoid occlusion factor), ParB, SMC and CcrZ (Veiga *et al*., 2011; Chan *et al*., 2020; Gallay *et al*., 2021).

Bacterial cell division also requires the activity of many associated components, often of ill-defined function. DivIVA and GpsB are two divisome proteins that are conserved within Firmicutes and the roles that they perform are well reviewed (Halbedel and Lewis, 2019; Hammond *et al*., 2019). DivIVA is a coiled-coil protein which binds to negatively curved membranes via its N-terminus, such as at the cell poles in rod shaped organisms and where the septum crosses the cell wall (Lenarcic *et al*., 2009; Ramamurthi and Losick, 2009). The N-terminal domain is linked to the C-terminal domain, via a short linker (Halbedel and Lewis, 2019), which facilitates oligomerisation into a tetramer (Muchová *et al*., 2002; Stahlberg *et al*., 2004; Rigden *et al*., 2008; Oliva *et al*., 2010). DivIVA localises to the site of division in *Bacillus subtilis* forming two rings around the Z-ring. This prevents the Min system from interacting with FtsZ, allowing division to continue, and stopping additional adjacent Z-rings from forming (Eswaramoorthy *et al*., 2011). DivIVA also interacts with the Spo0J/ParB system in *B. subtilis* and *Streptococcus pneumoniae* to facilitate chromosome segregation (Perry and Edwards, 2006; Fadda *et al*., 2007; Kloosterman *et al*., 2016). Little is known about the function of *S. aureus* DivIVA, with a null mutant having no significant changes to cellular morphology or division (Pinho and Errington, 2004). DivIVA is stabilised through interacting with DnaK, and plays a role in chromosome segregation, likely through an interaction with SMC (Bottomley *et al*., 2017).

GpsB is a homologue of DivIVA (Hammond *et al*., 2019). Both proteins have a similar overall structure, with a highly conserved N-terminal domain linked to a C-terminal domain (required for oligomerisation) via a short linker (Halbedel and Lewis, 2019). In *B. subtilis,* where it was first described, it was found to play a role in the switch between septal and peripheral peptidoglycan synthesis through interactions with PBP1 and MreC (Claessen *et al*., 2008; Tavares *et al*., 2008; Gamba *et al*., 2009). These observations suggest that GpsB acts as an adaptor protein to bring together different components of the divisome during the cell cycle (Cleverley *et al*., 2019). In *S. aureus* it has been shown that GpsB interacts with and bundles FtsZ filaments, stabilising the Z-ring and assisting with divisome recruitment (Eswara *et al*., 2018). GpsB also plays a role in linking cell division with wall teichoic acid display and synthesis (Hammond *et al*., 2022).

It has previously been reported that *gpsB* is essential (Eswara *et al*., 2018), and the function of *divIVA* remains relatively unknown in *S. aureus.* As *gpsB* is a homologue of *divIVA,* we aimed to find out if these genes have a collective or distinct roles in cell growth and division. In this study we utilised super resolution microscopy complemented with other molecular approaches to interrogate the function of *gpsB* and *divIVA* in *S. aureus.* We show that *gpsB* plays a role in cell shape determination.

## Results

### GpsB and DivIVA localise at the septum

As DiviVA and GpsB are paralogues they may be functionally related, and therefore share a localisation. Previous work has independently shown that DivIVA (Pinho and Errington, 2004) and GpsB (Eswara *et al*., 2018) localise to the *S. aureus* septum. To assess the co-localization of GpsB and DivIVA fusion constructs were produced using pOB (GpsB- mCherry) and pKASBAR (DivIVA-meGFP) and co-expressed in the same strain (SH1000 *gpsB::gpsB-mCherry kanR ΔdivIVA geh::divIVA-megfp*). Both GpsB-mCherry and DivIVA- meGFP localise at the septum when compared to a HADA label, which allows visualisation of new cell wall peptidoglycan synthesis by fluorescence microscopy (Kuru *et al*., 2012a) (Figure 1A). However, DivIVA-meGFP has a ‘dotty’ pattern of localization, which has previously been reported (Pinho and Errington, 2004), whereas GpsB-mCherry is seen to be forming smooth rings, suggesting these proteins are not co-localising. When viewing through the Z-stacks, it is clear that both GpsB-mCherry and DivIVA-meGFP are forming patterns at the developing septum (Video 1).

**Figure 1.**
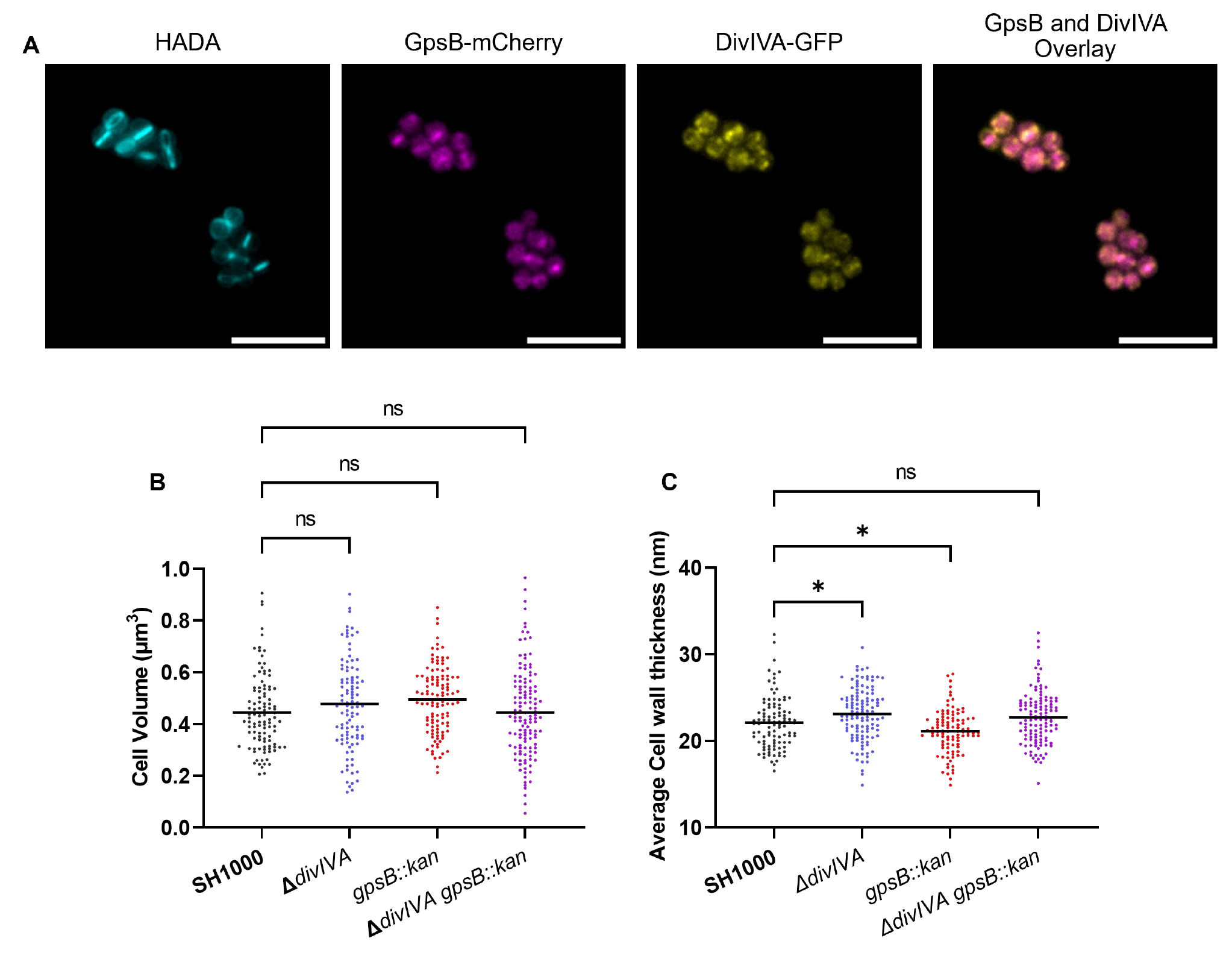
The localisation and role of DivIVA and GpsB. **(A)** Representative fluorescence microscopy images of SH1000 cells **(A)** labelled with HADA for 30 minutes and expressing GpsB-mCherry and DivIVA-GFP. An overlay of the localisations of GpsB-mCherry and DivIVA-GFP is also shown (scale bar represents 5 µm). **(B)** Cell volume analysis from SIM micrographs of SH1000 (black circles, n = 107 cells), SH1000 *ΔdivIVA* (blue circles, n = 104 cells), SH1000 *gpsB::kan* (red circles, n = 114 cells) and SH1000 *ΔdivIVA gpsB::kan* (purple circles, n = 130 cells). **(C)** Average cell wall thicknesses of SH1000 (black circles, n = 103 cells), SH1000 *ΔdivIVA* (blue circles, n = 120 cells, * *p* = 0.0283), SH1000 *gpsB::kan* (red circles, n = 101 cells, * *p* = 0.0401) and SH1000 *ΔdivIVA gpsB::kan* (purple circles, n = 119 cells). Results for **(B)** and **(C)** were analysed using a one-way ANOVA with multiple comparisons (ns *p* > 0.05).

### *gpsB* and *divIVA* mutations do not impact cell volume

Previous reports have suggested that GpsB is an essential protein in *S. aureus* (Santiago *et al*., 2015; Eswara *et al*., 2018), however here we were able to construct a marked deletion of the *gpsB* gene in SH1000 using piMay and thus it is non-essential in this background. A transposon inactivation *gpsB* mutant can also be found within the NARSA transposon library (Fey *et al*., 2013). A markerless *divIVA* deletion mutant was also constructed using pMAD, as well as a double mutant (SH1000 *gpsB::kan* Δ*divIVA*). No differences in growth were found between any of the mutants or when compared to the wild type (Supplementary Figure 1). Structured Illumination Microscopy (SIM) was used to analyse the cell volume of the mutants (Figure 1B and Supplementary Figure 2). No significant differences were observed in the volumes of the SH1000 Δ*divIVA*, SH1000 *gpsB::kan* or SH1000 Δ*divIVA gpsB::kan* strains compared to the wildtype SH1000. Next, TEM was used to interrogate the cell wall structure of these mutants (Figure 1C and Supplementary Figure 1). SH1000 Δ*divIVA* was found to have a slightly thicker peripheral cell wall than wildtype cells (*p =* 0.0283) while SH1000 *gpsB::kan* had a slightly thinner cell wall (*p =* 0.0401). No difference in cell wall thickness was observed in SH1000 Δ*divIVA gpsB::kan* compared to wildtype SH1000 (Figure 1C).

### Combinatorial mutagenesis reveals a role for *gpsB* and *divVIA* in cell volume determination

The roles of *gpsB* and *divIVA* in other species and the results of bacterial two-hybrid screens previously described in the literature (Steele *et al*., 2011; Bottomley *et al*., 2017) allowed us to determine a potential role for DivIVA and GpsB in various aspects of cell growth and division. As single and double mutants of *gpsB* and *divIVA* did not show any clear phenotype, further mutations, from the NARSA transposon library (Fey *et al*., 2013) (unless otherwise stated), were added in combination with SH1000 Δ*divIVA gpsB::kan* in order to provide information about DivIVA and GpsB.

### Teichoic Acids

*S. aureus* encodes three LytR-CpsA-Psr (LCP) proteins within its genome: *lcpA, lcpB* and *lcpC* (Over *et al*., 2011; Chan *et al*., 2013), all of which have a putative interaction with GpsB in a bacterial two-hybrid system (Kent, 2013). It has been proposed that the LCP family of proteins catalyse the transfer of WTA intermediates to the cell wall (Kawai *et al*., 2011; Chan *et al*., 2013). Severe phenotypic defects of *lcpA* mutants prevented establishment of multiple mutant strains (Over *et al*., 2011; Chan *et al*., 2013) and no significant difference could be found in cell volume when comparing SH1000 *lcpB::ery* and SH1000 *lcpB::ery* Δ*divIVA gpsB::kan* (Supplementary Figure 3A). An *lcpC* mutation, which has previously been shown to have the smallest impact on cell growth and morphology (Over *et al*., 2011; Chan *et al*., 2013), was also tested. SH1000 *lcpC::ery* was found to be significantly smaller than wildtype SH1000, while SH1000 *lcpC::ery* Δ*divIVA gpsB::kan* was significantly larger than SH1000 (Figure 2A). This increase in cell size was observed in all stages of the cell cycle (Supplementary Figure 4A). Strains SH1000 Δ*divIVA lcpC::ery* and SH1000 *gpsB::kan lcpC::ery* were produced to deconvolve these results. Both SH1000 Δ*divIVA lcpC::ery* and SH1000 *gpsB::kan lcpC::ery* showed no significant differences in cell volume to SH1000 (Figure 2C and D). This suggests that the phenotype is present for SH1000 Δ*divIVA lcpC::ery* and SH1000 *gpsB::kan lcpC::ery* but is exacerbated further in a triple mutant. SH1000 Δ*divIVA gpsB::kan lcpC::ery* was complemented using a pKASBAR plasmid, containing native *gpsB,* which integrates into the *geh* locus of *S. aureus*. A control of an empty pKASBAR was also used. As expected, both SH1000 Δ*divIVA gpsB::kan lcpC::ery* and SH1000 Δ*divIVA gpsB::kan lcpC::ery geh::pKASBAR* were significantly larger than wildtype SH1000, whereas SH1000 Δ*divIVA gpsB::kan lcpC::ery geh::gpsB* was the same size (Supplementary Figure 5A).

**Figure 2.**
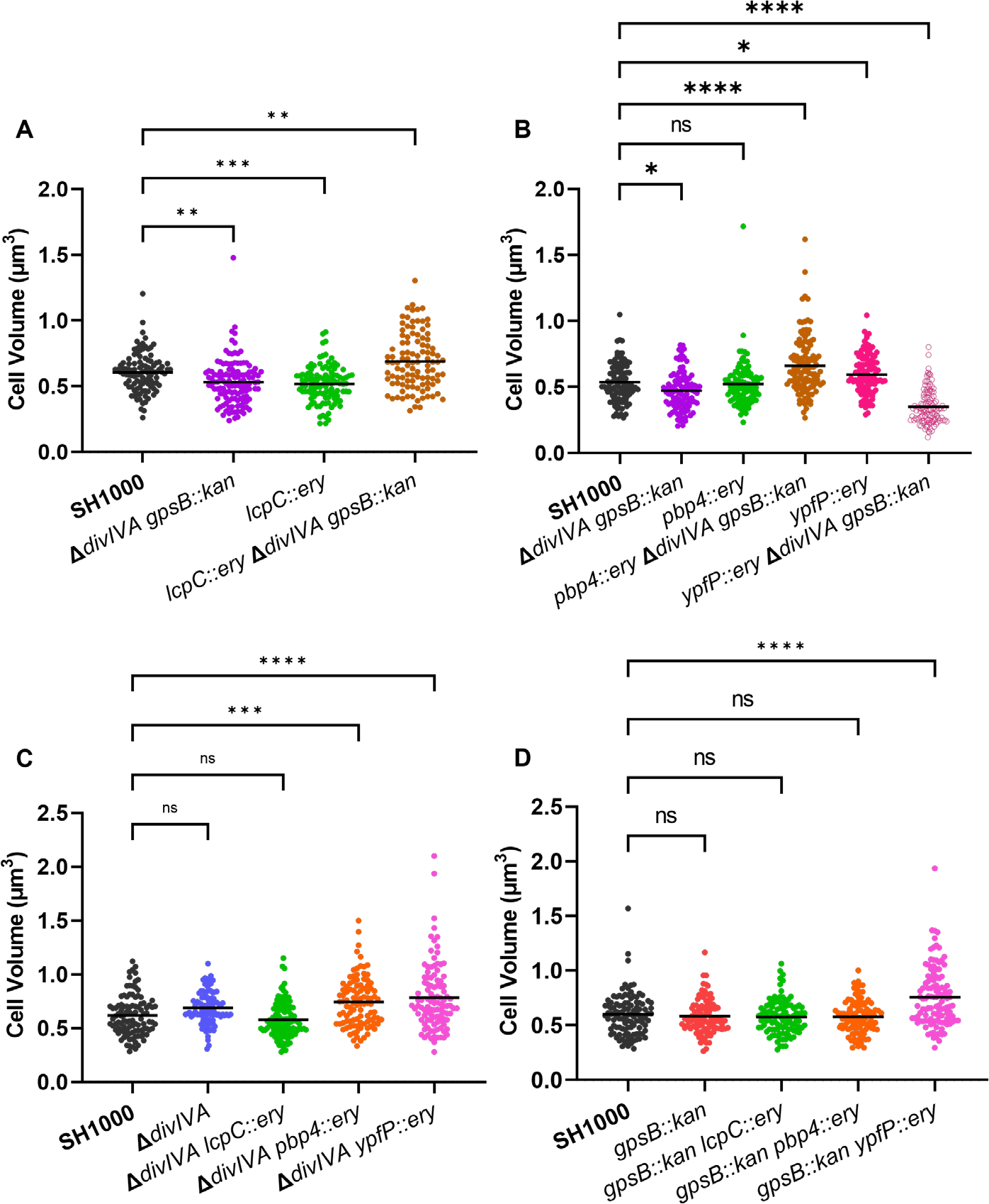
Functional interaction between DivIVA and GpsB with other components. **(A)** Cell volume analysis from SIM micrographs of SH1000 (black circles, n = 104 cells), SH1000 *ΔdivIVA gpsB::kan* (purple circles, n = 124 cells, *p* = 0.0028), SH1000 *lcpC::ery* (green circles, n = 107 cells, *p* = 0.0004), SH1000 *ΔdivIVA gpsB::kan lcpC::ery* (brown circles, n = 104 cells, *p* = 0.0014). **(B)** Cell volume analysis from SIM micrographs of SH1000 (black circles, n = 105 cells), SH1000 *ΔdivIVA gpsB::kan* (purple circles, n = 107 cells, *p* = 0.0192), SH1000 *pbp4::ery* (green circles, n = 102 cells, *p* = 0.9711), SH1000 *ΔdivIVA gpsB::kan pbp4::ery* (brown circles, n = 125 cells, *p* < 0.0001), SH1000 *ypfP::ery* pink circles, n = 113 cells, *p* = 00348), SH1000 *ΔdivIVA gpsB::kan ypfP::ery* open pink circles, n = 117 cells, *p* < 0.0001). **(C)** Cell volume analysis from SIM micrographs of SH1000 (black circles, n = 104 cells), SH1000 *ΔdivIVA* (blue circles, n = 102 cells, = = 0.0673), SH1000 *ΔdivIVA lcpC::ery* (green circles, n = 119 cells, *p* = 0.4148), SH1000 *ΔdivIVA pbp4::ery* (orange circles, n = 101 cells, *p* = 0.0002), SH1000 *ΔdivIVA ypfP::ery* (pink circles, n = 105 cells, *p* < 0.0001). **(D)** Cell volume analysis from SIM micrographs of SH1000 (black circles, n = 108 cells), SH1000 *gpsB::kan* (red circles, n = 100 cells, 0.9257), SH1000 *gpsB::kan lcpC::ery* (green circles, n = 103 cells, *p* = 0.7417), SH1000 *gpsB::kan pbp4::ery* (orange circles, n = 104 cells, *p* = 0.8137), SH1000 *gpsB::kan ypfP::ery* (pink circles, n = 114 cells, *p* < 0.0001). Results analysed using a one-way ANOVA with multiple comparisons.

TarO, the first enzyme in the biosynthetic pathway of wall teichoic acids (Soldo *et al*., 2002; Atilano *et al*., 2010), was shown to interact with both DivIVA and GpsB (Kent, 2013). Although SH1000 *tarO::ery* (Salamaga *et al*., 2021) and SH1000 *tarO::ery* Δ*divIVA gpsB::kan* both show a significantly increased cell volume compared to SH1000, *ΔdivIVA* and *gpsB::kan* did not alter this phenotype (Supplementary Figure 4B).

Due to the apparent interaction between DivIVA and GpsB and proteins involved in WTA synthesis and display, a potential role for LTA, which has been shown to be involved in cell division (Gründling and Schneewind, 2007), was also investigated. As the synthesis of LTA is essential via the action of LtaS (Gründling and Schneewind, 2007), *ypfP,* which has a 87% reduction in LTA content (Fedtke *et al*., 2007), was investigated. SH1000 *ypfP::ery* has a significantly greater volume than SH1000, while SH1000 *ypfP::ery* Δ*divIVA gpsB::kan* was significantly smaller than wildtype (Figure 2B), suggesting that LTA may be important in DivIVA or GpsB function. The decrease in cell volume was observed in all stages of the cell cycle (Supplementary Figure 4C). SH1000 Δ*divIVA ypfP::ery* and SH1000 *gpsB::kan ypfP::ery* were constructed to further analyse this phenotype. Both SH1000 Δ*divIVA ypfP::ery* and SH1000 *gpsB::kan ypfP::ery* show an increase in cell volume compared to SH1000 (Figure 2C and D), the same phenotype as the SH1000 *ypfP::ery*. Therefore, the loss of *divIVA, gpsB* and *ypfP* are all required for the reduced volume phenotype of the triple mutant. Complementation of SH1000 Δ*divIVA gpsB::kan ypfP::ery* with pKASBAR expressing native *gpsB* restored the increased cell volume phenotype (Supplementary Figure 5).

Both DivIVA and GpsB interact with PBP4 (Kent, 2013). SH1000 *pbp4::ery* had no significant difference in cell volume compared to parental SH1000, whereas SH1000 *pbp4::ery* Δ*divIVA gpsB::kan* has a significantly greater volume than SH1000 (Figure 2B). This increase in cell volume was only seen in cells with no or an incomplete septum (Supplementary Figure 4B). The results were deconvolved by constructing and analysing SH1000 Δ*divIVA pbp4::ery* and SH1000 *gpsB::kan pbp4::ery*. SH1000 Δ*divIVA pbp4::ery* shows a significantly greater increase in cell volume compared to SH1000 (Figure 2C). However, SH1000 *gpsB::kan pbp4::ery* shows no significant difference in cell volume compared to wildtype (Figure 2D), demonstrating that only a loss of *divIVA* is required for the phenotype. This result was complemented using pKASBAR expressing native *divIVA*. (Supplementary Figure 5B).

### Chromosome Segregation

Previous research has shown a link between DivIVA and the segregation of chromosomes prior to cell division (Bottomley *et al*., 2017). The nucleoid occlusion protein Noc prevents the septa of a dividing cell to form over a chromosome, acting as an important checkpoint for chromosome segregation (Veiga *et al*., 2011). No chromosome segregation phenotype could be found for SH1000 *noc::ery* Δ*divIVA gpsB::kan,* based on DAPI staining, and no differences in cell volume were observed (Supplementary Figure 3).

DivIVA of *Corynebacterium glutamicum* binds to ParB and helps to orient the chromosome for cell division, as well as resulting in mobilisation of DivIVA (Giacomelli *et al*., 2022). While SH1000 *parB::ery* was slightly but significantly larger than SH1000, SH1000 *parB::ery* Δ*divIVA gpsB::kan* showed no significant volume differences to SH1000 or SH1000 Δ*divIVA gpsB::kan* (Supplementary Figure 3D), and no abnormalities in chromosome segregation were observed.

### YpsA

Previous work has shown that *gpsB* is in a syntenous relationship with *ypsA* within the genomes of Firmicutes, including *S. aureus* (Brzozowski *et al*., 2019), and conservation of such organisation often indicates shared function (Aravind, 2000; Huynen *et al*., 2000). Due to *gpsB* being encoded directly downstream from *ypsA,* we were unable to transduce both *ypsA::ery* and *gpsB::kan* into a single strain, so instead analysed SH1000 *ypsA::ery ΔdivIVA.* No differences could be found in cell volume (Supplementary Figure 3E) or chromosome segregation for SH1000 *ypsA::ery ΔdivIVA*.

### GpsB plays a role in *S. aureus* cell circularity

*S. aureus* has previously been shown to elongate during the cell cycle (Monteiro *et al*., 2015), resulting in a long-axis and a short-axis. Calculating the ratio between these two axes allows the extent of elongation to be calculated as previously reported (Reichmann *et al*., 2019), and acts as a measure of circularity. In this study, we calculated the ratio by dividing the short axis (axis perpendicular to the long axis) by the long axis (axis perpendicular to the septum). Using the short/long axis ratio, a value of 1 indicates that the cell is perfectly circular, whereas the smaller the ratio the more elongated the cell is.

The short/long axis ratio was calculated for Δ*divIVA* and *gpsB::kan* mutants (Figure 3A). SH1000 Δ*divIVA* had no significant difference in short/long axis ratio compared to SH1000, while both SH1000 *gpsB::kan* and SH1000 Δ*divIVA gpsB::kan* had a significantly greater ratio, meaning that the cells are more circular, or less elongated, than wildtype cells. Both SH1000 *gpsB::kan* and SH1000 Δ*divIVA gpsB::kan* have a significantly higher short/long axis ratios with incomplete septa compared to SH1000 and SH1000 Δ*divIVA*, but there is no difference with no septa (Figure 3B). Complementation with *gpsB* being expressed from the *geh* locus using pKASBAR (Bottomley *et al*., 2014), restored the elongation phenotype, with no difference in short/long axis ratio between the wildtype and complemented strains (Figure 3C and D).

**Figure 3.**
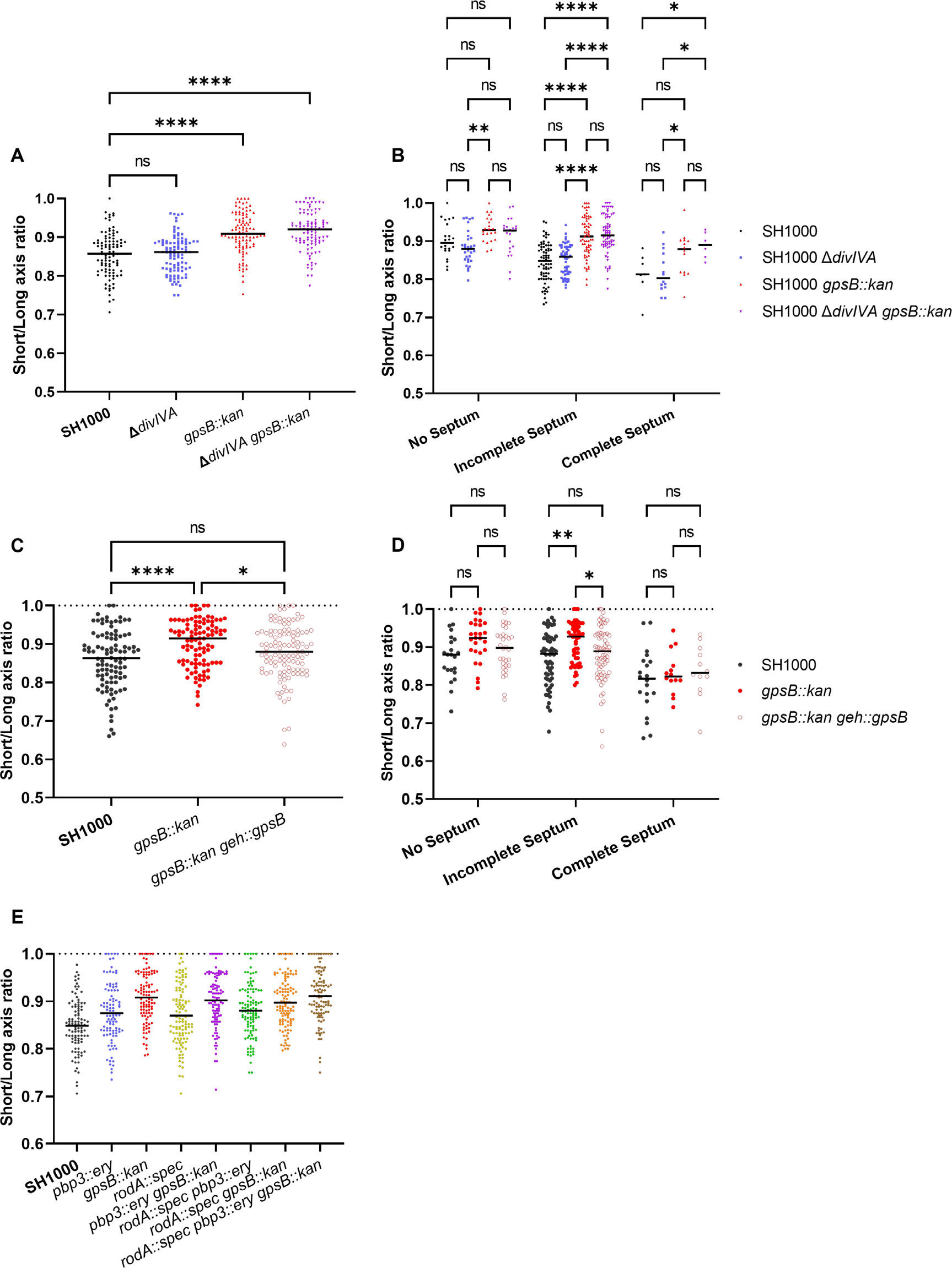
GpsB has a role in cell circularity. **(A)** The short/long axis ratio of SH1000 (black circles, n = 100 cells), SH1000 *ΔdivIVA* (blue circles, n = 100 cells, *p* > 0.9999), SH1000 *gpsB::kan* (red circles, n = 101 cells, *p* < 0.0001) and SH1000 *ΔdivIVA gpsB::kan* (purple circles, n = 101 cells, *p* < 0.0001). Data was analysed using a one-way ANOVA. **(B)** The short/long axis ratios from **(A)** organised by the stage of the cell cycle cells were in (*p* values ** *p* = 0.0024, **** *p* < 0.0001). Analysed using a two-way ANOVA with multiple comparisons. **(C)** The short/long axis ratio of SH1000 (black circles, n = 100 cells), SH1000 *gpsB::kan* (red circles, n = 101 cells), SH1000 *gpsB::kan geh::gpsB* (open red circles, n = 113 cells). Analysed using a one-way ANOVA with multiple comparisons (ns *p* = 0.0903, * *p* = 0.0456, **** *p* < 0.00001). **(D)** The short/long axis ratios from **(C)** organised by the stage of the cell cycle cells were in (*p* values * *p* = 0.0201, ** *p* = 0.0024). Analysed using a two-way ANOVA with multiple comparisons. **(E)** The short/long axis ratio of SH1000 (black circles, n = 100 cells), SH1000 *pbp3::ery* (blue circles, n = 99 cells), SH1000 *gpsB::kan* (red circles, n = 102 cells), SH1000 *rodA::spec* (yellow circles, n = 112 cells), SH1000 *pbp3::ery gpsB::kan* (purple circles, n = 107 cells), SH1000 *pbp3::ery rodA::spec* (green circles, n = 108 cells), SH1000 *rodA::spec gpsB::kan* (orange circles, n = 111 cells) and SH1000 *rodA::spec pbp3::ery gpsB::kan* (brown circles, n = 102 cells). The *p* values for **(E)** are in supplementary table 4.

It has previously been reported that RodA and PBP3 form a cognate pair that is important for peripheral PG synthesis and the elongation of *S. aureus* (Reichmann *et al*., 2019). All single, double and triple mutant permutations were created for *rodA, gpsB* and *pbp3* and compared (Figure 3E). SH1000 *ppb3::ery* and SH1000 *rodA::spec pbp3::ery* were significantly more circular than SH1000. SH1000 *gpsB::ery* was found to be significantly more circular than SH1000, SH1000 *rodA::spec,* SH1000 *pbp3::ery* and SH1000 *rodA::spec pbp3::ery*. SH1000 *gpsB::kan* showed no differences in circularity to to SH1000 *rodA::spec pbp3::ery gpsB::kan* (Figure 3E and Supplementary Table 4).

### GpsB influences PG synthesis and the localisation of PBPs

The localisation of PG synthesis was determined by measuring the fluorescence ratio (FR) at the septum and the periphery of cells that have been sequentially labelled with ADA-DA and HADA (Figure 4A) for 5 mins each to follow septal development (Tinajero-Trejo *et al*., 2022). The higher the FR, the greater the PG synthesis at the septum compared to the periphery. SH1000 *gpsB::kan* had a significantly higher FR than wildtype SH1000 (Figure 4B).

**Figure 4.**
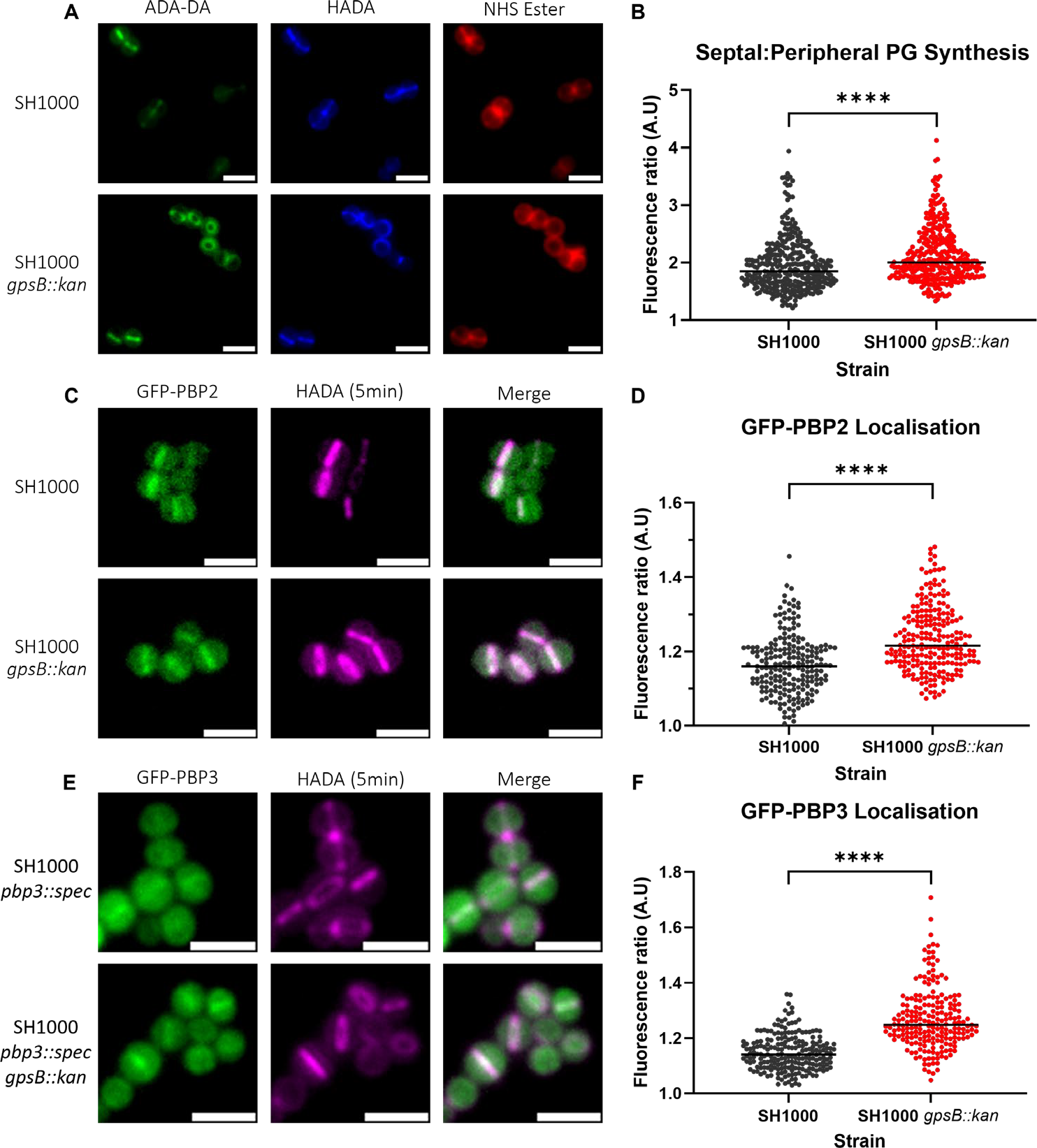
The role of GpsB in PG synthesis and PBP localisation. **(A)** Representative micrographs showing ADA-DA (labelled with Atto488, green), HADA (blue) and NHS Ester 555 (red) labelled cells of SH1000 and SH1000 *gpsB::kan* (scale bar represents 2 µm)**. (B) F**luorescence ratio (FR) values for the septal/peripheral ADA-DA signal in SH1000 (black circles, n = 322 cells) and SH1000 *gpsB::kan* (red circles, n = 312 cells). **(C)** Representative images of micrographs showing the localization of GFP-PBP2 in SH1000 pLOW-GFP-PBP2 and SH1000 *gpsB::kan* pLOW-PBP2-gfp (scale bar represents 2 µm). **(D)** Fluorescence ratio values for the GFP-PBP2 signal in SH1000 (black circles, n = 209) and SH1000 *gpsB::kan* (red circles, n = 229). Representative images of micrographs showing the localization of GFP-PBP3 in **(E)** SH1000 *pbp3::spec* pLOW*-gfp-pbp3* and SH1000 *pbp3::spec gpsB::kan* pLOW-*gfp-pbp3* (scale bar represents 2 µm) **(F)** Fluorescence ratio values for the GFP-PBP3 signal in SH1000 *pbp3::spec* pLOW*-gfp-pbp3* (black circles, n = 211) and SH1000 *pbp3::spec gpsB::kan* pLOW-*gfp-pbp3* (red circles, n = 212). All were compared using a Mann-Whitney test (**** *p* < 0.0001).

As PG synthesis is altered in SH1000 *gpsB::kan*, we determined PBP localisation at the septum compared to the cell periphery using PBP fluorescent reporter fusions. PBP2 is the major PG transpeptidase in *S. aureus* responsible for the bulk of PG synthesis (Pinho *et al*., 2001b, 2001a). Imaging SH1000 and SH1000 *gpsB::kan* expressing GFP-PBP2 (Figure 4C) showed a significant increase in FR for SH1000 *gpsB::kan* (Figure 4D), indicating that there is more GFP-PBP2 in the septum of dividing cells.

PBP3 plays a role in the elongation of *S. aureus* due to off-septal PG synthesis (Reichmann *et al*., 2019). A GFP-PBP3 construct was made in pLOW under the control of the *Ppcn* promoter, which was then transduced into SH1000 *pbp3::spec* and SH1000 *pbp3::spec gpsB::kan* so the only expressed copy of PBP3 was GFP-PBP3. To test if the fusion was functional, the circularity of SH1000 (Supplementary Figure 6A), SH1000 *pbp3::spec* (Supplementary Figure 6B), SH1000 *pbp3::spec* pLOW-*gfp-pbp3* (Supplementary Figure 6C) and SH1000 *pbp3::spec gpsB::kan* pLOW-*gfp-pbp3* (Supplementary Figure 6D) were compared. SH1000 *pbp3::spec* is significantly more circular than SH1000. SH1000 *pbp3::spec* pLOW-*gfp-pbp3* was found to be significantly less circular than SH1000 *pbp3::spec* and have no difference to SH1000 but could not complement the increased circularity of *gpsB::kan* (Supplementary Figure 6E). SH1000 *pbp3::spec gpsB::kan* pLOW- *gfp-pbp3* (Figure 4E) had a significantly higher FR for GFP-PBP3 than SH1000 *pbp3::spec* pLOW-*gfp-pbp3* (Figure 4F), demonstrating increased PBP3 at the septum in the absence of GpsB.

## Discussion

DivIVA and GpsB are paralogues in *S. aureus,* suggesting that they may have overlapping roles. The two proteins were not found to co-localise however, and through screening we noted interesting phenotypes for SH1000 *ΔdivIVA pbp4::ery*, SH1000 *ΔdivIVA gpsB::kan ypfP::ery* and SH1000 *ΔdivIVA gpsB::kan lcpC::ery*. A change in morphology was also noted for *gpsB::kan* mutants. SH1000 *gpsB::kan* was found to be more circular, or less elongated, than wildtype cells, a phenotype that was more pronounced than for a PBP3 mutant (Reichmann *et al*., 2019). This phenotype is due to an increased proportion of PG synthesis at the septum of *gpsB::kan* cells, which was associated with an increased proportion of PBP2 and PBP3 localising at the septum of SH1000 *gpsB::kan.* The results presented here suggest that GpsB plays a role in *S. aureus* PG synthesis regulation, specifically in regulating between septal and peripheral synthesis.

Both *pbp4* and *lcpC* combination knockouts were shown to produce larger cells (Figure 2). The activity PBP4 is known to be regulated by WTA, with WTA being responsible for the localisation of PBP4 at the septum and determining the level of PG crosslinking via the activity of PBP4 (Atilano *et al*., 2010). LcpC is involved in the ligation and display of WTA on PG after its synthesis. As DivIVA and GpsB both interact with TarO (Kent, 2013), an apparent link between DivIVA and GpsB with WTA is implicated. It has recently been demonstrated that GpsB directly interacts with TarG, involved in the export of WTA and found to localise with the divisome complex (Hammond *et al*., 2022). GpsB, and now also DivIVA, play a role in linking WTA and cell division together, perhaps for the regulation of division. The increase in cell volume observed with these combination mutants could be due to the deregulation of PG synthases, potentially due to a disconnect with WTA (Atilano *et al*., 2010). While previously having been shown to play a role of regulation with WTA, we have now also suggested a link with DivIVA, GpsB and LTA, despite GpsB not being a part of the *S. aureus* LTA synthesis machinery complex (Reichmann *et al*., 2014). LTAs play a role in septal placement as well as determining morphology, as cells lacking *ypfP* were misshapen (Kiriukhin *et al*., 2001; Gründling and Schneewind, 2007; Oku *et al*., 2009). A *ypfP* mutant was shown to be larger than SH1000, which is previously noted (Reichmann *et al*., 2014). However, cells that lacked *ypfP, divIVA* and *gpsB* were significantly smaller, suggesting a deregulation of cell growth, widening the molecules important for the function of both DivIVA and GpsB from WTA to teichoic acids in general. It has been suggested that GpsB acts as an adapter protein bringing together components of the diviosome to regulate division (Cleverley *et al*., 2019). Our results suggesting a link between DivIVA, GpsB, WTA and LTA further add evidence for this hypothesis.

Due to the changes in volume being observed for the *pbp4, lcpC* and *ypfP* combination mutants, it is possible that a change in PG synthesis is responsible for this. Interestingly, DivIVA has been shown to interact with PBP1 (Bottomley *et al*., 2017). With DivIVA having an interaction with DivIC (which itself also interacts with WTA) it is possible that they act in similar pathways. DivIC has been found to play a role in septal formation and in the regulation and localisation of PBP2 (Tinajero-Trejo *et al*., 2022). With DivIVA having a wide range of interactions, it is conceivable that DivIVA helps to regulate similar processes and brings together components of the divisome. The lack of a strong phenotype associated with loss of *divIVA* implies that its function is redundant in the cell, or assisting the activity of other proteins, perhaps through stabilising their interactions.

Previous studies have shown that during the cell cycle *S. aureus* elongates slightly (Monteiro *et al*., 2015) and this elongation is due, in part, to the activity of RodA and PBP3 incorporating PG to the side wall (Reichmann *et al*., 2019). While GpsB has been shown to only directly interact with PBP4 (Steele *et al*., 2011; Kent, 2013), it was found to be important for the localisation of PBP2 and PBP3. As the localization of these PBPs are altered in a *gpsB* mutant, GpsB may occlude accumulation of these PBPs at the septum, allowing insertion of PG into the peripheral cell wall and subsequent elongation of the cell. Regulation of septal and peripheral PG synthesis has previously been described in *B. subtilis*, where direct interaction between PBP1 and MreC switched the cell between the two modalities of synthesis (Claessen *et al*., 2008; Tavares *et al*., 2008; Gamba *et al*., 2009). GpsB plays a similar role in the rod-shaped *Listeria monocytogenes,* switching synthesis between the periphery and the septum by interacting with PBPA1 (Cleverley *et al*., 2016; Rismondo *et al*., 2017). In the ovococcoid *S. pneumoniae,* GpsB also regulates peripheral cell wall and septal synthesis. GpsB was shown to activate PBP2a and localise PBP2x to the late stage septum, with a model suggesting GpsB inhibited cell elongation by restricting the activity of PBP2b (Rued *et al*., 2017).

GpsB has been shown to interact with and stabilise FtsZ, concentrating its GTPase activity and helping to activate treadmilling for cytokinesis (Eswara *et al*., 2018). The transition between stages of cell division must be tightly controlled to prevent the chromosome being damaged during segregation and ensuring the two daughter cells are identical (Lund *et al*., 2018; Saraiva *et al*., 2020). As GpsB stabilises Z-ring formation, and then helps to activate treadmilling, as well as regulating PG synthesis at the septum and periphery, it could be that GpsB is regulating and bringing together components to temporally regulate cell division, ensuring that stages are occurring at the right time and order. In *B. subtilis*, DiviVA is known to localise to negatively curved membranes via its N-terminal lipid binding domain (Halbedel and Lewis, 2019). Such sites include the emerging division site due to the constriction of the membrane by FtsZ (Harry and Lewis, 2003; Eswaramoorthy *et al*., 2011), where it sandwiches the Z-ring by forming a ring on either side (Eswaramoorthy *et al*., 2011).

Here we have furthered the knowledge of *S. aureus* DivIVA and GpsB function, showing a link between their roles and teichoic acids. Also, we have shown a role for GpsB in cell shape determination. We propose a model whereby GpsB plays a role in the elongation of cells by blocking binding of PBPs (specifically 2 and 3) at the septum so that a greater proportion of PG synthesis occurs at the cell periphery, resulting in elongation.

## Materials and Methods

### Bacterial growth conditions, plasmids and oligonucleotides

The list of strains, plasmids and oligonucleotides used in this work are listed in Supplementary Tables 1, 2 and 3 respectively. All strains were cultured at 37°C with shaking at 200 rpm. For *S. aureus,* mid-exponential phase was defined as an OD600 of 0.4–0.8. *E. coli* was cultured in Luria-Bertani (LB) broth or agar plus 100 μg/ml ampicillin. *S. aureus* strains were grown in tryptic soy broth (TSB) (Bacto) or agar (TSA). Where required antibiotics were added at the following concentrations: 5 μg/ml erythromycin (Ery) 25 μg/ml lincomycin (Lin), 10 μg/ml chloramphenicol (Cm), 5 μg/ml tetracycline (Tet), 50 μg/ml kanamycin (Kan) and 100 μg/ml spectinomycin (Spec).

For growth curves overnight *S. aureus* cultures were adjusted to OD600 0.05 in TSB and incubated at 37°C with shaking at 200 rpm to grow for 8 h. Samples were taken every hour and OD600 was measured. Direct cell counts were also performed by serial dilution in PBS and plating onto TSA. The number of colony forming units (CFU) were directly counted after incubation. Growth curves were performed in triplicate. *E. coli* transformations and DNA manipulations were performed according to previously described methodology (Sambrook and Russell, 2001).

### Construction of *S. aureus* mutants

Unless otherwise stated, all vectors were constructed in *E. coli* NEB5α (New England Biolabs) following previously described methods (Gibson *et al*., 2009; Lund *et al*., 2018) before passage through *S. aureus* RN4220 for DNA methylation (Novick and Morse, 1967). Finally, constructs were transduced into *S. aureus* SH1000 using phage Φ11. Transductions and transformations were confirmed by PCR. Genomic DNA of SH1000 was used as a template for *S. aureus* gene amplification. Genomic DNA was isolated by incubating *S*. *aureus* cells in 2.5 μg/ml lysostaphin prior to extraction using Qiagen DNeasy Blood and Tissue kit (Cat no. 69506) in accordance with the manufacturer’s instructions.

### NARSA Library

*S*. *aureus* transposon mutants were obtained from the NARSA library (Bae *et al*., 2008; Fey *et al*., 2013). Transposons were transduced from the library to the recipient strain and confirmed by PCR.

### SH1000 *gpsB::kan*

To delete native *gpsB*, fragments encompassing 1000 bp upstream and downstream of *gpsB* were amplified using oligos piMAY*_gpsB_up_F/R* and piMAY*_gpsB_down_F/R*. A kanamycin resistance cassette was amplified from pGL433 (Wheeler *et al*., 2015) using oligos *pGL433_kan_F/R* and included in between the upstream and downstream fragments to allow selection of deletion mutants. The products were ligated into piMAY cut with KpnI and NotI and fragments were combined using Gibson assembly creating piMAY *gpsB-ko*. The plasmid was electroporated into RN4220 at 30 °C. The plasmid integrated through a single-crossover event at 37 °C and the chromosomal DNA fragment containing the deletion cassette was transduced into SH1000 to produce SJF4925. Colonies were selected based on kanamycin resistance and tetracycline sensitivity.

### SH1000 *ΔdivIVA – pMAD*

To construct pMAD Δ*divIVA,* 1000bp upstream and downstream of *divIVA* was amplified using oligo pairs pMAD_*divIVA_1/2* and *pMAD*_*divIVA_3/4.* pMAD was cut with BglII and EcoRI. The fragments were combined using Gibson assembly producing pMAD-Δ*divIVA*. This construct was transformed into RN4220, and a single crossover event occurred. The integrated pMAD-Δ*divIVA* was transduced into SH1000 where pMAD was excised by double crossover as previously described (Arnaud *et al*., 2004). This produced strain SJF4814.

### SH1000 *gpsB::gpsB-mCherry*

A C-terminal fusion of GpsB with mCherry was designed in pOB (Horsburgh *et al*., 2002) and synthesised by GENEWIZ UK Ltd. The synthesised plasmid was directly transformed into RN4220 (where it integrated into the chromosome through a single crossover event) and transduced into SH1000 (selecting for strains that were kan^R^ but sensitive to ery) to produce strain SJF5643.

### SH1000 *ΔdivIVA geh::divIVA-GFP*

A C-terminal fusion of DivIVA with GFP was created within pKASBAR *tet* (Bottomley *et al*., 2014). The insert region was synthesised by GENEWIZ UK Ltd and amplified using oligos pKB*-divIVA-F/-R*. The amplified fragment was then ligated into pKASBAR *tet* cut with BamHI and EcoRI using Gibson assembly (creating pKASBAR*-divIVA-gfp*). The plasmid was transduced into RN4220 with integration at the *geh* locus being confirmed by disruption of lipase production on Baird-Parker medium and PCR. The chromosomal fragment was then transduced into SJF4814 (SH1000 *ΔdivIVA*) to produce strain SJF5299.

### SH1000 *gpsB::gpsB-mCherry kanR ΔdivIVA geh::divIVA-megfp*

The chromosomal *gpsB::gpsB-mCherry* was transduced into SJF5299 and confirmed by PCR to produce SJF5669.

### SH1000 *ΔdivIVA geh::divIVA*

The *divIVA* locus including the native promoter (178 bp upstream) within pKASBAR *tet*. DNA fragments were made using oligos pKASBAR*_divIVA_F/R*. The resulting fragment was ligated into pKASBAR *tet* cut with BamHI and EcoRI using Gibson assembly creating pKASBAR-*divIVA*. The construct was electroporated into RN4220, with integration at the *geh* locus being confirmed by disruption of lipase production on Baird-Parker medium and PCR. The chromosomal fragment containing integrated pKASBAR-*divIVA* was transduced into SJF4814 to produce SJF4899.

### SH1000 *gpsB::kan geh::gpsB*

The *gpsB* locus including the native promoter within pKASBAR *tet*. DNA fragments were made using oligos pKASBAR*_gpsB_F/R.* The resulting fragment was ligated into pKASBAR *tet* cut with BamHI and EcoRI using Gibson assembly creating pKASBAR-*gpsB*. The construct was electroporated into RN4220, with integration at the *geh* locus being confirmed by disruption of lipase production on Baird-Parker medium and PCR. The chromosomal fragment containing integrated pKASBAR-*gpsB* was transduced into SJF4925 to produce SJF4956.

### SH1000 *pbp3::spec pLOW-Ppcn-gfp-pbp3*

An N-terminal fusion of PBP3 with GFP was synthesised and cloned into pLOW under the control of a *Ppcn* promoter by GENEWIZ UK Ltd. This plasmid was electroporated into RN4220 and then transduced into SH1000 *pbp3::spec* and SH1000 *pbp3::spec gpsB::kan* to produce SJF5950 and SJF5951 respectively.

### Transmission electron microscopy (TEM)

TEM was performed as in (Sutton *et al*., 2021). Briefly, samples were fixed overnight in 2.5 % (w/v) glutaraldehyde at 4 °C. Samples were washed in PBS and secondary fixation was performed with 2 % (w/v) osmium tetroxide for 2 hours. After washing, samples were dehydrated in incrementally increasing concentrations of ethanol and then incubated in propylene oxide. Samples were infiltrated overnight in a 50% (v/v) propylene oxide to 50% (v/v) Epon resin mixture overnight, which was then replaced with pure Epon resin for 4 hours, which was then replaced with fresh resin for another 4 hours. Polymerization was then performed in fresh resin at 60 °C for 48–72 hours. 80 nm thin sections were taken and stained with 3% (w/v) aqueous uranyl acetate followed by Reynold’s lead citrate. Sections were imaged using a FEI Tecnai T12 Spirit Transmission Electron Microscope operating at 80 kV. Images were recorded using a Gatan Orius SC1000B bottom mounted CCD camera. TEM images were analysed using Fiji software (Schindelin *et al*., 2012). Cell wall thickness was measured as previously described (Sutton *et al*., 2021).

### Labelling of strains for fluorescence microscopy

*S. aureus* strains were grown overnight in TSB (with appropriate antibiotics) which were used to inoculate fresh TSB to an OD600 of 0.05. Cells were then grown to mid exponential phase (OD600 of ∼0.5) before being labelled. Samples were protected from light throughout staining until imaged.

### Fluorescent D-amino acids (FDAA)

500 µM of 7-hydroxycoumarin-3-carboxylic acid-amino-D-alanine (HADA) or 1mM of azido-Dalanyl-D-alanine (ADA-DA) (Kuru *et al*., 2012b; Monteiro *et al*., 2015; Lund *et al*., 2018) was added to cells for 5 or 30 min and incubated at 37 °C with shaking. Cells were then washed twice in PBS at 4 °C. The azide group of ADA-DA was labelled (post-fixation) with 5 µg ml^−1^ Alexa Fluor 488 Alkyne using the Click-iT™ Cell Reaction Buffer Kit (Invitrogen) according to the manufacturer instructions.

### NHS Ester

Cells were resuspended in PBS and incubated at 4 °C for 5 mins with 8 µg ml^−1^ Alexa Fluor 555 NHS ester (Invitrogen). Cells were then washed in PBS.

### Fixation by paraformaldehyde

After labelling cells were incubated with 2% (w/v) paraformaldehyde (PFA) for 30 min at room temperature and then washed twice in water.

### DAPI

After fixation cells were resuspended in water containing 2 µg/ml DAPI (Sigma) for 5 mins at room temperature on a rotary shaker. Samples were then washed twice in water before mounting.

### Widefield Epifluorescence microscopy

Cells were mounted onto poly-L-Lysine coated slides (Sigma) using SlowFade^TM^ Gold antifade reagent (Thermo Fisher) and then imaged using a Nikon Ti inverted microscope fitted with a Lumencor Spectra X light engine. Images were obtained with a 100x PlanApo (1.4 NA) oil objective 1.518 RI oil and an Andora Zyla sCMOS camera was used for detection.

### OMX microscopy (Structured Illumination Microscopy (SIM))

SIM was performed as previously described (Lund *et al*., 2018). Briefly, coverslips (High-precision, 1.5H, 22 ± 22 mm, 170 ± 5 mm, Marienfeld) were sonicated in 1 M KOH for 15 min before being washed and then incubated in poly-L-Lysine solution for 30 min. Coverslips were then washed and dried before fixed cells (suspended in water) were dried onto coverslips and mounted with SlowFade^TM^ Gold antifade reagent (Thermo Fisher).

SIM was performed using a v4 DeltaVision OMX 3D-SIM system fitted with a Blaze module (Applied Precision, GE Healthcare, Issaquah, USA) with lasers used to illuminate samples. For each Z-slice (0.125 nm), images were taken for in five phase shifts and three angles. To reconstruct images the software Softworx (GE Healthcare) was used with optimisation for a 1.516 immersion oil. The same software was used for deconvolution and image alignment.

### Microscopy analysis

All measurements from microscopy images were made using Fiji (Schindelin *et al*., 2012). Unless otherwise stated micrographs presented are maximum intensity projections of Z-stacks.

### Cell volume analysis

Cell volume analysis from widefield microscopy and SIM were analysed as previously described (Zhou *et al*., 2015). Long and short axis measurements were taken for each cell and the volume was calculated using the equation for volume of a prolate spheroid:

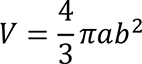

### Cell elongation short/long cell axis ratio

Axis ratio was adapted from previously published methodology (Reichmann *et al*., 2019). To calculate the short/long axis cell ratio as an indicator of cell shape the short axis measurement of each cell (as taken for volume) was divided by the long axis to give a ratio where cells are perfectly circular at 1.0 and more elongated as the value decreases.

### Fluorescence ratio septal/peripheral

Fluorescence ratio (FR) was calculated as previously published (Tinajero-Trejo *et al*., 2022). FR was calculated using fluorescence at the septum of cells with an incomplete septum with fluorescence measured between the cell periphery and the annulus. This was divided by the mean fluorescence at the lateral cell walls.

### Statistical analysis

Statistical analysis was performed using Prism version 9.31 (GraphPad).

## Acknowledgments

This paper is dedicated to Mark Cooke, who sadly passed away before the project was completed. This work was funded by the Wellcome Trust (212197/Z/19/Z). We gratefully acknowledge the Wolfson Light Microscopy facility for their support and assistance in this work. We are grateful to Lucia Lafage, Laia Pasquina-Lemonche and Chris Hill for help and advice. J.A.F.S would like to thank Amy Tooke, Oliver Carnell, Josie Pyrah, Eric Pollitt and Chloe Allen for encouragement throughout the project.

## References

Aravind, L. (2000). Guilt by association: contextual information in genome analysis. Genome Res 10, 1074–1077. doi: 10.1101/gr.10.8.1074.

Arnaud, M., Chastanet, A., and Débarbouillé, M. (2004). New vector for efficient allelic replacement in naturally nontransformable, low-GC-content, gram-positive bacteria. Appl Environ Microbiol 70, 6887–6891. doi: 10.1128/AEM.70.11.6887-6891.2004.

Atilano, M. L., Pereira, P. M., Yates, J., Reed, P., Veiga, H., Pinho, M. G., et al. (2010). Teichoic acids are temporal and spatial regulators of peptidoglycan cross-linking in *Staphylococcus aureus*. PNAS 107, 18991–18996. doi: 10.1073/pnas.1004304107.

Bae, T., Glass, E. M., Schneewind, O., and Missiakas, D. (2008). Generating a collection of insertion mutations in the *Staphylococcus aureus* genome using bursa aurealis. Methods Mol Biol 416, 103–116. doi: 10.1007/978-1-59745-321-9_7.

Bottomley, A. L., Kabli, A. F., Hurd, A. F., Turner, R. D., Garcia-Lara, J., and Foster, S. J. (2014). *Staphylococcus aureus* DivIB is a peptidoglycan-binding protein that is required for a morphological checkpoint in cell division. Molecular Microbiology 94, 1041–1064. doi: 10.1111/mmi.12813.

Bottomley, A. L., Liew, A. T. F., Kusuma, K. D., Peterson, E., Seidel, L., Foster, S. J., et al. (2017). Coordination of Chromosome Segregation and Cell Division in *Staphylococcus aureus*. Front Microbiol 8, 1575. doi: 10.3389/fmicb.2017.01575.

Brzozowski, R. S., Huber, M., Burroughs, A. M., Graham, G., Walker, M., Alva, S. S., et al. (2019). Deciphering the Role of a SLOG Superfamily Protein YpsA in Gram-Positive Bacteria. Front Microbiol 10, 623. doi: 10.3389/fmicb.2019.00623.

Cabeen, M. T., and Jacobs-Wagner, C. (2005). Bacterial cell shape. Nat Rev Microbiol 3, 601–610. doi: 10.1038/nrmicro1205.

Chan, H., Söderström, B., and Skoglund, U. (2020). Spo0J and SMC are required for normal chromosome segregation in *Staphylococcus aureus*. Microbiologyopen 9, e999. doi: 10.1002/mbo3.999.

Chan, Y. G. Y., Frankel, M. B., Dengler, V., Schneewind, O., and Missiakas, D. (2013). *Staphylococcus aureus* Mutants Lacking the LytR-CpsA-Psr Family of Enzymes Release Cell Wall Teichoic Acids into the Extracellular Medium. J Bacteriol 195, 4650–4659. doi: 10.1128/JB.00544-13.

Claessen, D., Emmins, R., Hamoen, L. W., Daniel, R. A., Errington, J., and Edwards, D. H. (2008). Control of the cell elongation-division cycle by shuttling of PBP1 protein in Bacillus subtilis. Mol Microbiol 68, 1029–1046. doi: 10.1111/j.1365-2958.2008.06210.x.

Cleverley, R. M., Rismondo, J., Lockhart-Cairns, M. P., Van Bentum, P. T., Egan, A. J. F., Vollmer, W., et al. (2016). Subunit Arrangement in GpsB, a Regulator of Cell Wall Biosynthesis. Microb Drug Resist 22, 446–460. doi: 10.1089/mdr.2016.0050.

Cleverley, R. M., Rutter, Z. J., Rismondo, J., Corona, F., Tsui, H.-C. T., Alatawi, F. A., et al. (2019). The cell cycle regulator GpsB functions as cytosolic adaptor for multiple cell wall enzymes. Nat Commun 10, 261. doi: 10.1038/s41467-018-08056-2.

Eswara, P. J., Brzozowski, R. S., Viola, M. G., Graham, G., Spanoudis, C., Trebino, C., et al. (2018). An essential *Staphylococcus aureus* cell division protein directly regulates FtsZ dynamics. eLife 7, e38856. doi: 10.7554/eLife.38856.

Eswaramoorthy, P., Erb, M. L., Gregory, J. A., Silverman, J., Pogliano, K., Pogliano, J., et al. (2011). Cellular Architecture Mediates DivIVA Ultrastructure and Regulates Min Activity in Bacillus subtilis. mBio 2, e00257–11. doi: 10.1128/mBio.00257-11.

Fadda, D., Santona, A., D’Ulisse, V., Ghelardini, P., Ennas, M. G., Whalen, M. B., et al. (2007). Streptococcus pneumoniae DivIVA: Localization and Interactions in a MinCD- Free Context. Journal of Bacteriology 189, 1288–1298. doi: 10.1128/JB.01168-06.

Fedtke, I., Mader, D., Kohler, T., Moll, H., Nicholson, G., Biswas, R., et al. (2007). A *Staphylococcus aureus* ypfP mutant with strongly reduced lipoteichoic acid (LTA) content: LTA governs bacterial surface properties and autolysin activity. Mol Microbiol 65, 1078–1091. doi: 10.1111/j.1365-2958.2007.05854.x.

Fey, P. D., Endres, J. L., Yajjala, V. K., Widhelm, T. J., Boissy, R. J., Bose, J. L., et al. (2013). A genetic resource for rapid and comprehensive phenotype screening of nonessential *Staphylococcus aureus* genes. mBio 4, e00537–00512. doi: 10.1128/mBio.00537-12.

Gallay, C., Sanselicio, S., Anderson, M. E., Soh, Y. M., Liu, X., Stamsås, G. A., et al. (2021). CcrZ is a pneumococcal spatiotemporal cell cycle regulator that interacts with FtsZ and controls DNA replication by modulating the activity of DnaA. Nat Microbiol 6, 1175–1187. doi: 10.1038/s41564-021-00949-1.

Gamba, P., Veening, J.-W., Saunders, N. J., Hamoen, L. W., and Daniel, R. A. (2009). Two-step assembly dynamics of the Bacillus subtilis divisome. J Bacteriol 191, 4186– 4194. doi: 10.1128/JB.01758-08.

Giacomelli, G., Feddersen, H., Peng, F., Martins, G. B., Grafemeyer, M., Meyer, F., et al. (2022). Subcellular Dynamics of a Conserved Bacterial Polar Scaffold Protein. Genes (Basel) 13, 278. doi: 10.3390/genes13020278.

Gibson, D. G., Young, L., Chuang, R.-Y., Venter, J. C., Hutchison, C. A., and Smith, H. O. (2009). Enzymatic assembly of DNA molecules up to several hundred kilobases. Nat Methods 6, 343–345. doi: 10.1038/nmeth.1318.

Gründling, A., and Schneewind, O. (2007). Synthesis of glycerol phosphate lipoteichoic acid in *Staphylococcus aureus*. Proc Natl Acad Sci U S A 104, 8478–8483. doi: 10.1073/pnas.0701821104.

Halbedel, S., and Lewis, R. J. (2019). Structural basis for interaction of DivIVA/GpsB proteins with their ligands. Molecular Microbiology 111, 1404–1415. doi: 10.1111/mmi.14244.

Hammond, L. R., Sacco, M. D., Khan, S. J., Spanoudis, C., Hough-Neidig, A., Chen, Y., et al. (2022). GpsB Coordinates Cell Division and Cell Surface Decoration by Wall Teichoic Acids in *Staphylococcus aureus*. Microbiol Spectr 10, e0141322. doi: 10.1128/spectrum.01413-22.

Hammond, L. R., White, M. L., and Eswara, P. J. (2019). ¡vIVA la DivIVA! *J Bact*e*riol* 201, e00245–19. doi: 10.1128/JB.00245-19.

Harry, E. J., and Lewis, P. J. (2003). Early targeting of Min proteins to the cell poles in germinated spores of Bacillus subtilis: evidence for division apparatus-independent recruitment of Min proteins to the division site. Molecular Microbiology 47, 37–48. doi: 10.1046/j.1365-2958.2003.03253.x.

Hartman, B. J., and Tomasz, A. (1984). Low-affinity penicillin-binding protein associated with beta-lactam resistance in *Staphylococcus aureus*. Journal of Bacteriology 158, 513–516. doi: 10.1128/jb.158.2.513-516.1984.

Horsburgh, M. J., Wharton, S. J., Cox, A. G., Ingham, E., Peacock, S., and Foster, S. J. (2002). MntR modulates expression of the PerR regulon and superoxide resistance in *Staphylococcus aureus* through control of manganese uptake. Mol Microbiol 44, 1269–1286. doi: 10.1046/j.1365-2958.2002.02944.x.

Huynen, M., Snel, B., Lathe, W., and Bork, P. (2000). Predicting Protein Function by Genomic Context: Quantitative Evaluation and Qualitative Inferences. Genome Res 10, 1204–1210.

Kawai, Y., Marles-Wright, J., Cleverley, R. M., Emmins, R., Ishikawa, S., Kuwano, M., et al. (2011). A widespread family of bacterial cell wall assembly proteins. EMBO J 30, 4931–4941. doi: 10.1038/emboj.2011.358.

Kent, V. (2013). Cell wall architecture and the role of wall teichoic acid in *Staphylococcus aureus*. Available at: https://etheses.whiterose.ac.uk/5599/ [Accessed April 19, 2023].

Kiriukhin, M. Y., Debabov, D. V., Shinabarger, D. L., and Neuhaus, F. C. (2001). Biosynthesis of the Glycolipid Anchor in Lipoteichoic Acid of *Staphylococcus aureus* RN4220: Role of YpfP, the Diglucosyldiacylglycerol Synthase. J Bacteriol 183, 3506– 3514. doi: 10.1128/JB.183.11.3506-3514.2001.

Kloosterman, T. G., Lenarcic, R., Willis, C. R., Roberts, D. M., Hamoen, L. W., Errington, J., et al. (2016). Complex polar machinery required for proper chromosome segregation in vegetative and sporulating cells of Bacillus subtilis. Molecular Microbiology 101, 333–350. doi: 10.1111/mmi.13393.

Kuru, E., Hughes, H. V., Brown, P. J., Hall, E., Tekkam, S., Cava, F., et al. (2012a). In Situ probing of newly synthesized peptidoglycan in live bacteria with fluorescent D-amino acids. Angew Chem Int Ed Engl 51, 12519–12523. doi: 10.1002/anie.201206749.

Kuru, E., Hughes, H. V., Brown, P. J., Hall, E., Tekkam, S., Cava, F., et al. (2012b). In Situ probing of newly synthesized peptidoglycan in live bacteria with fluorescent D-amino acids. Angew Chem Int Ed Engl 51, 12519–12523. doi: 10.1002/anie.201206749.

Lenarcic, R., Halbedel, S., Visser, L., Shaw, M., Wu, L. J., Errington, J., et al. (2009). Localisation of DivIVA by targeting to negatively curved membranes. EMBO J 28, 2272–2282. doi: 10.1038/emboj.2009.129.

Loskill, P., Pereira, P. M., Jung, P., Bischoff, M., Herrmann, M., Pinho, M. G., et al. (2014). Reduction of the peptidoglycan crosslinking causes a decrease in stiffness of the *Staphylococcus aureus* cell envelope. Biophys J 107, 1082–1089. doi: 10.1016/j.bpj.2014.07.029.

Lund, V. A., Wacnik, K., Turner, R. D., Cotterell, B. E., Walther, C. G., Fenn, S. J., et al. (2018). Molecular coordination of *Staphylococcus aureus* cell division. eLife 7, e32057. doi: 10.7554/eLife.32057.

Monteiro, J. M., Fernandes, P. B., Vaz, F., Pereira, A. R., Tavares, A. C., Ferreira, M. T., et al. (2015). Cell shape dynamics during the staphylococcal cell cycle. Nat Commun 6, 8055. doi: 10.1038/ncomms9055.

Muchová, K. N., Kutejová, E., Scott, D. J., Brannigan, J. A., Lewis, R. J., Wilkinson, A. J., et al. (2002). Oligomerization of the Bacillus subtilis division protein DivIVA. Microbiology (Reading) 148, 807–813. doi: 10.1099/00221287-148-3-807.

Novick, R. P., and Morse, S. I. (1967). IN VIVO TRANSMISSION OF DRUG RESISTANCE FACTORS BETWEEN STRAINS OF *STAPHYLOCOCCUS AUREUS*. J Exp Med 125, 45–59.

Oku, Y., Kurokawa, K., Matsuo, M., Yamada, S., Lee, B.-L., and Sekimizu, K. (2009). Pleiotropic Roles of Polyglycerolphosphate Synthase of Lipoteichoic Acid in Growth of *Staphylococcus aureus* Cells. J Bacteriol 191, 141–151. doi: 10.1128/JB.01221-08.

Oliva, M. A., Halbedel, S., Freund, S. M., Dutow, P., Leonard, T. A., Veprintsev, D. B., et al. (2010). Features critical for membrane binding revealed by DivIVA crystal structure. EMBO J 29, 1988–2001. doi: 10.1038/emboj.2010.99.

Over, B., Heusser, R., McCallum, N., Schulthess, B., Kupferschmied, P., Gaiani, J. M., et al. (2011). LytR-CpsA-Psr proteins in *Staphylococcus aureus* display partial functional redundancy and the deletion of all three severely impairs septum placement and cell separation. FEMS Microbiology Letters 320, 142–151. doi: 10.1111/j.1574-6968.2011.02303.x.

Perry, S. E., and Edwards, D. H. (2006). The Bacillus subtilis DivIVA Protein Has a Sporulation-Specific Proximity to Spo0J. Journal of Bacteriology 188, 6039–6043. doi: 10.1128/JB.01750-05.

Pinho, M. G., de Lencastre, H., and Tomasz, A. (2000). Cloning, characterization, and inactivation of the gene pbpC, encoding penicillin-binding protein 3 of *Staphylococcus aureus*. J Bacteriol 182, 1074–1079. doi: 10.1128/JB.182.4.1074-1079.2000.

Pinho, M. G., de Lencastre, H., and Tomasz, A. (2001a). An acquired and a native penicillin-binding protein cooperate in building the cell wall of drug-resistant staphylococci. Proc Natl Acad Sci U S A 98, 10886–10891. doi: 10.1073/pnas.191260798.

Pinho, M. G., and Errington, J. (2004). A divIVA null mutant of *Staphylococcus aureus* undergoes normal cell division. FEMS Microbiol Lett 240, 145–149. doi: 10.1016/j.femsle.2004.09.038.

Pinho, M. G., and Errington, J. (2005). Recruitment of penicillin-binding protein PBP2 to the division site of *Staphylococcus aureus* is dependent on its transpeptidation substrates. Mol Microbiol 55, 799–807. doi: 10.1111/j.1365-2958.2004.04420.x.

Pinho, M. G., Filipe, S. R., de Lencastre, H., and Tomasz, A. (2001b). Complementation of the essential peptidoglycan transpeptidase function of penicillin-binding protein 2 (PBP2) by the drug resistance protein PBP2A in *Staphylococcus aureus*. J Bacteriol 183, 6525–6531. doi: 10.1128/JB.183.22.6525-6531.2001.

Pinho, M. G., Kjos, M., and Veening, J.-W. (2013). How to get (a)round: mechanisms controlling growth and division of coccoid bacteria. Nat Rev Microbiol 11, 601–614. doi: 10.1038/nrmicro3088.

Ramamurthi, K. S., and Losick, R. (2009). Negative membrane curvature as a cue for subcellular localization of a bacterial protein. Proc Natl Acad Sci U S A 106, 13541– 13545. doi: 10.1073/pnas.0906851106.

Reichmann, N. T., Piçarra Cassona, C., Monteiro, J. M., Bottomley, A. L., Corrigan, R. M., Foster, S. J., et al. (2014). Differential localization of LTA synthesis proteins and their interaction with the cell division machinery in *Staphylococcus aureus*. Mol Microbiol 92, 273–286. doi: 10.1111/mmi.12551.

Reichmann, N. T., Tavares, A. C., Saraiva, B. M., Jousselin, A., Reed, P., Pereira, A. R., et al. (2019). SEDS-bPBP pairs direct lateral and septal peptidoglycan synthesis in *Staphylococcus aureus*. Nat Microbiol 4, 1368–1377. doi: 10.1038/s41564-019-0437-2.

Rigden, M. D., Baier, C., Ramirez-Arcos, S., Liao, M., Wang, M., and Dillon, J.-A. R. (2008). Identification of the coiled-coil domains of Enterococcus faecalis DivIVA that mediate oligomerization and their importance for biological function. J Biochem 144, 63–76. doi: 10.1093/jb/mvn044.

Rismondo, J., Bender, J. K., and Halbedel, S. (2017). Suppressor Mutations Linking gpsB with the First Committed Step of Peptidoglycan Biosynthesis in Listeria monocytogenes. J Bacteriol 199, e00393–16. doi: 10.1128/JB.00393-16.

Rued, B. E., Zheng, J. J., Mura, A., Tsui, H.-C. T., Boersma, M. J., Mazny, J. L., et al. (2017). Suppression and synthetic-lethal genetic relationships of ΔgpsB mutations indicate that GpsB mediates protein phosphorylation and penicillin-binding protein interactions in Streptococcus pneumoniae D39. Molecular Microbiology 103, 931– 957. doi: 10.1111/mmi.13613.

Salamaga, B., Kong, L., Pasquina-Lemonche, L., Lafage, L., von Und Zur Muhlen, M., Gibson, J. F., et al. (2021). Demonstration of the role of cell wall homeostasis in *Staphylococcus aureus* growth and the action of bactericidal antibiotics. Proc Natl Acad Sci U S A 118, e2106022118. doi: 10.1073/pnas.2106022118.

Sambrook, J., and Russell, D. W. (2001). Molecular Cloning: A Laboratory Manual. 3rd ed. Cold Spring Harbor, NY: Cold Spring Harbor Laboratory Press.

Santiago, M., Matano, L. M., Moussa, S. H., Gilmore, M. S., Walker, S., and Meredith, T. C. (2015). A new platform for ultra-high density *Staphylococcus aureus* transposon libraries. BMC Genomics 16, 252. doi: 10.1186/s12864-015-1361-3.

Saraiva, B. M., Sorg, M., Pereira, A. R., Ferreira, M. J., Caulat, L. C., Reichmann, N. T., et al. (2020). Reassessment of the distinctive geometry of *Staphylococcus aureus* cell division. Nat Commun 11, 4097. doi: 10.1038/s41467-020-17940-9.

Schindelin, J., Arganda-Carreras, I., Frise, E., Kaynig, V., Longair, M., Pietzsch, T., et al. (2012). Fiji: an open-source platform for biological-image analysis. Nat Methods 9, 676–682. doi: 10.1038/nmeth.2019.

Soldo, B., Lazarevic, V., and Karamata, D. 2002 (2002). tagO is involved in the synthesis of all anionic cell-wall polymers in Bacillus subtilis 168aaThe EMBL accession number for the nucleotide sequence reported in this paper is AJ004803. Microbiology 148, 2079–2087. doi: 10.1099/00221287-148-7-2079.

Stahlberg, H., Kutejová, E., Muchová, K., Gregorini, M., Lustig, A., Müller, S. A., et al. (2004). Oligomeric structure of the Bacillus subtilis cell division protein DivIVA determined by transmission electron microscopy. Mol Microbiol 52, 1281–1290. doi: 10.1111/j.1365-2958.2004.04074.x.

Steele, V. R., Bottomley, A. L., Garcia-Lara, J., Kasturiarachchi, J., and Foster, S. J. (2011). Multiple essential roles for EzrA in cell division of *Staphylococcus aureus*. Molecular Microbiology 80, 542–555. doi: 10.1111/j.1365-2958.2011.07591.x.

Sutton, J. A. F., Carnell, O. T., Lafage, L., Gray, J., Biboy, J., Gibson, J. F., et al. (2021). *Staphylococcus aureus* cell wall structure and dynamics during host-pathogen interaction. PLOS Pathogens 17, e1009468. doi: 10.1371/journal.ppat.1009468.

Szwedziak, P., Wang, Q., Bharat, T. A. M., Tsim, M., and Löwe, J. (2014). Architecture of the ring formed by the tubulin homologue FtsZ in bacterial cell division. Elife 3, e04601. doi: 10.7554/eLife.04601.

Tavares, J. R., de Souza, R. F., Meira, G. L. S., and Gueiros-Filho, F. J. (2008). Cytological characterization of YpsB, a novel component of the Bacillus subtilis divisome. J Bacteriol 190, 7096–7107. doi: 10.1128/JB.00064-08.

Tinajero-Trejo, M., Carnell, O., Kabli, A. F., Pasquina-Lemonche, L., Lafage, L., Han, A., et al. (2022). The *Staphylococcus aureus* cell division protein, DivIC, interacts with the cell wall and controls its biosynthesis. Commun Biol 5, 1–13. doi: 10.1038/s42003-022-04161-7.

Turner, R. D., Vollmer, W., and Foster, S. J. (2014). Different walls for rods and balls: the diversity of peptidoglycan. Mol Microbiol 91, 862–874. doi: 10.1111/mmi.12513.

Typas, A., Banzhaf, M., Gross, C. A., and Vollmer, W. (2011). From the regulation of peptidoglycan synthesis to bacterial growth and morphology. Nat Rev Microbiol 10, 123–136. doi: 10.1038/nrmicro2677.

Veiga, H., Jorge, A. M., and Pinho, M. G. (2011). Absence of nucleoid occlusion effector Noc impairs formation of orthogonal FtsZ rings during *Staphylococcus aureus* cell division. Molecular Microbiology 80, 1366–1380. doi: 10.1111/j.1365-2958.2011.07651.x.

Vollmer, W., Blanot, D., and De Pedro, M. A. (2008). Peptidoglycan structure and architecture. FEMS Microbiology Reviews 32, 149–167. doi: 10.1111/j.1574-6976.2007.00094.x.

Wacnik, K., Rao, V. A., Chen, X., Lafage, L., Pazos, M., Booth, S., et al. (2022). Penicillin-Binding Protein 1 (PBP1) of *Staphylococcus aureus* Has Multiple Essential Functions in Cell Division. mBio 0, e00669–22. doi: 10.1128/mbio.00669-22.

Wheeler, R., Turner, R. D., Bailey, R. G., Salamaga, B., Mesnage, S., Mohamad, S. A. S., et al. (2015). Bacterial Cell Enlargement Requires Control of Cell Wall Stiffness Mediated by Peptidoglycan Hydrolases. mBio 6, e00660. doi: 10.1128/mBio.00660-15.

Wyke, A. W., Ward, J. B., Hayes, M. V., and Curtis, N. A. (1981). A role in vivo for penicillin-binding protein-4 of *Staphylococcus aureus*. Eur J Biochem 119, 389–393. doi: 10.1111/j.1432-1033.1981.tb05620.x.

Zhou, X., Halladin, D. K., Rojas, E. R., Koslover, E. F., Lee, T. K., Huang, K. C., et al. (2015). Mechanical crack propagation drives millisecond daughter cell separation in *Staphylococcus aureus*. Science 348, 574–578. doi: 10.1126/science.aaa1511.

