## Supplementary Data for "The roles of GpsB and DivIVA in *Staphylococcus aureus* growth and division"

| Name | Genotype | Source |
|---|---|---|
| ***E. coli*** | | |
| NEB5α | | New England Biolabs |
| ***Staphylococcus aureus*** | | |
| SH1000 | Functional *rsbU*+ derivative of 8325–4 | (Horsburgh et al., 2002) |
| SJF4925 | SH1000 *gpsB::kan* | This Study |
| SJF4814 | SH1000 Δ*divIVA* | This Study |
| SJF5093 | SH1000 Δ*divIVA gpsB::kan* | This Study |
| SJF5640 | RN4220 *gpsB::gpsB-mCherry kan* | This Study |
| SJF5643 | SH1000 *gpsB::gpsB-mCherry kan* | This Study |
| SJF5299 | SH1000 *ΔdivIVA geh::divIVA-gfp* | This Study |
| SJF5669 | SH1000 *ΔdivIVA geh::divIVA-gfp gpsB::gpsB-mCherry kan* | This Study |
| SJF4899 | SH1000 *ΔdivIVA geh::divIVA* | This Study |
| SJF4956 | SH1000 *gpsB::kan geh::gpsB* | This Study |
| NE415 | JE2 *lcpC::Tn(ery)* | (Fey et al., 2013) |
| SJF5448 | SH1000 *lcpC::ery* | This Study |
| SJF5623 | SH1000 *gpsB::kan lcpC::ery* | This Study |
| SJF5620 | SH1000 Δ*divIVA lcpC::ery* | This Study |
| SJF5449 | SH1000 Δ*divIVA gpsB::kan lcpC::ery* | This Study |
| SJF5743 | SH1000 Δ*divIVA gpsB::kan lcpC::ery geh::pKASBAR* | This Study |
| SJF5744 | SH1000 Δ*divIVA gpsB::kan lcpC::ery geh::gpsB* | This Study |
| SJF4425 | SH1000 *pbp4::ery* | (Sutton et al., 2021) |
| SJF5624 | SH1000 *gpsB::kan pbp4::ery* | This Study |
| SJF5621 | SH1000 Δ*divIVA pbp4::ery* | This Study |
| SJF5492 | SH1000 Δ*divIVA gpsB::kan pbp4::ery* | This Study |
| SJF5747 | SH1000 Δ*divIVA pbp4::ery geh::pKASBAR* | This Study |
| SJF5748 | SH1000 Δ*divIVA pbp4::ery geh::divIVA* | This Study |
| NE1663 | JE2 *ypfP::Tn(ery)* | (Fey et al., 2013) |
| SJF5526 | SH1000 *ypfP::ery* | This Study |
| SJF5625 | SH1000 *gpsB::kan ypfP::ery* | This Study |
| SJF5622 | SH1000 Δ*divIVA ypfP::ery* | This Study |
| SJF5527 | SH1000 Δ*divIVA gpsB::kan ypfP::ery* | This Study |
| SJF5745 | SH1000 Δ*divIVA gpsB::kan ypfP::ery geh::pKASBAR* | This Study |
| SJF5746 | SH1000 Δ*divIVA gpsB::kan ypfP::ery geh::gpsB* | This Study |
| NE486 | JE2 *noc::Tn(ery)* | (Fey et al., 2013) |
| SJF5442 | SH1000 *noc::ery* | This Study |
| SJF5443 | SH1000 Δ*divIVA gpsB::kan noc::ery* | This Study |
| NE1778 | JE2 *lcpB::Tn(ery)* | (Fey et al., 2013) |

| | | |
|---|---|---|
| SJF5446 | SH1000 *lcpB::ery* | This Study |
| SJF5447 | SH1000 Δ*divIVA gpsB::kan lcpB::ery* | This Study |
| NE1697 | JE2 *ypsA::Tn(ery)* | (Fey et al., 2013) |
| SJF5469 | SH1000 *ypsA::ery* | This Study |
| SJF5470 | SH1000 Δ*divIVA gpsB::kan ypsA::ery* | This Study |
| NE772 | JE2 *parB::Tn(ery)* | (Fey et al., 2013) |
| SJF5444 | SH1000 *parB::ery* | This Study |
| SJF5445 | SH1000 Δ*divIVA gpsB::kan parB::ery* | This Study |
| SJF5289 | SH1000 *tarO::ery* | (Salamaga et al., 2021) |
| SJF5482 | SH1000 Δ*divIVA gpsB::kan tarO::ery* | This Study |
| SJF4956 | SH1000 *gpsB::kan geh::gpsB* | This Study |
| SJF4421 | SH1000 *pbp3*::*ery* | (Wacnik et al., 2022) |
| SJF4422 | SH1000 *pbp3*::*spec* | (Wacnik et al., 2022) |
| NE1598 | JE2 *rodA::Tn(ery)* | (Fey et al., 2013) |
| SJF4837 | SH1000 *rodA::spec* | This Study |
| SJF5596 | SH1000 *pbp3*::*ery gpsB::kan* | This Study |
| SJF5579 | SH1000 *rodA::spec pbp3*::*ery* | This Study |
| SJF5578 | SH1000 *rodA::spec gpsB::kan* | This Study |
| SJF5597 | SH1000 *rodA::spec pbp3*::*ery gpsB::kan* | This Study |
| SJF5659 | SH1000 *pLOW-ppcn-gfp-pbp2* | (Tinajero-Trejo et al., 2022) |
| SJF5671 | SH1000 *gpsB::kan pLOW-Ppcn-gfp-pbp2* | This Study |
| SJF5750 | SH1000 *pbp3::spec pLOW-Ppcn-gfp-pbp3* | This Study |
| SJF5751 | SH1000 *pbp3::spec gpsB::kan* pLOW-*Ppcn-gfp-pbp3* | This Study |
| RN4220 | Restriction deficient transformation recipient | (Kreiswirth et al., 1983) |
| Ery, erythromycin resistance; Tet, tetracycline resistance; Kan, kanamycin resistance; Spec, Spectinomycin. | | |

**Supplementary Table 1 Strains used in this study**

| Name | Relevant genotype/markers | Source |
|---|---|---|
| pKASBAR *ery* | pUC18 containing *attP* and an *ery* resistance cassette. Amp, Ery | (Bottomley et al., 2014) |
| pKASBAR *tet* | pUC18 containing *attP* and a *tet* resistance cassette. Amp, Tet | (Bottomley et al., 2014) |
| pKASBAR-*divIVA* | pKASBAR *tet* encoding the whole of *divIVA* with 178 bp upstream to include native promoter and ribosome binding site. Fragment synthesised and cloned into BamHI and EcoRI sites | This Study |
| pKASBAR-*gpsB* | pKASBAR *tet* encoding the whole of *gpsB* with 1170 bp upstream to include native promoter and ribosome binding site. Fragment synthesised and cloned into BamHI and BglII sites. | This Study |
| pKASBAR-*divIVA-gfp* | pKASBAR *tet* encoding 1000bp upstream of *divIVA,* followed by the whole of *divIVA*, *linker B* and *gfp*. Insert was synthesised by Genewiz, amplified using *pKB-divIVA-F/-R* primers, and cloned into pKASBAR *tet* cut with BamHI and EcoRI | This Study |
| pOB-*gpsB-mCherry* | pOB carrying *gpsB* with the stop codon removed and a *linkerA-mCherry* fusion attached followed by a $kan^R$ cassette. Plasmid also has 658bp upstream of *gpsB* and 1000bp downstream to allow recombination. Ery, Amp, kan | GENEWIZ UK Ltd |
| piMAY | Temperature-sensitive Gram-positive replicon from pVE6007 with an *E. coli* replicon with tetracycline resistance cassette; Tet, Cm | (Monk et al., 2012) |
| piMAY *gpsB-ko* | piMAY carrying 1000bp upstream and 1000bp downstream *gpsB* with a kanamycin cassette between the two sequences. piMAY was cut with KpnI NotI. Tet, Cm, Kan | This Study |
| pMAD | *E. coli-S. aureus* shuttle vector with temperature-sensitive origin of replication in *S. aureus* and promoterless bgaB; Amp, Ery | (Arnaud et al., 2004) |
| pMAD *divIVA-ko* | pMAD carrying 1000bp upstream and 1000bp downstream of *divIVA.* pMAD was cut with BglII and EcoRI; Ery, Amp | This Study |
| pGL433 | Vector carrying *kan* cassette suitable for selection in Gram-positive bacteria; Kan | (Wheeler et al., 2015) |

| | | |
|---|---|---|
| pLOW | SK41-type low copy number plasmid; Amp, Ery | (Liew et al., 2011) |
| pLOW-*gfp-pbp2* | pLOW expressing a *gfp-pbp2* fusion under control of the penicillinase constitutive promoter (Ppcn); Ery | (Tinajero-Trejo et al., 2022) |
| pLOW-*gfp-pbp3* | pLOW expressing a *gfp-pbp3* fusion under control of the penicillinase constitutive promoter (Ppcn); Ery | Genewiz UK LTD This Study |
| Amp, ampicillin resistance; Ery, erythromycin resistance; Tet, tetracycline resistance; Kan, kanamycin resistance; Cm, chloramphenicol resistance. | | |

**Supplementary Table 2 Plasmids used in this study**

| **Oligonucleotide name (and restriction site)** | **Sequence (5’ to 3’)*** | **Use** | **Source** |
|---|---|---|---|
| *gpsB-up-F* | atttctataaaagctacgtcactgtg | Amplifies the *gpsB* locus of *S. aureus* | This study |
| *gpsB-down-R* | tttcatacgtcgtatcaaggtc | | This study |
| *divIVA-up-F* | tgcaacagttagttctttaaggttag | Amplifies the *divIVA* locus of *S. aureus* | This study |
| *divIVA-down-R* | aggcattaataacgtttctttgtaatc | | This study |
| *parB_F* | cgaacccgtagacacctcat | Amplifies the *parB* locus of *S. aureus* | This study |
| *parB_R* | agcgattttagttgcaatgt | | This study |
| *ypfP_F* | aaactaacggagggtggcta | Amplifies the *ypfP* locus of *S. aureus* | This study |
| *ypfP_R* | gcaatggatgtaactgttggc | | This study |
| *RodA_FWD* | aaatctatagctgatatcactg | Amplifies the *rodA* locus of *S. aureus* | |
| *RodA_REV* | ttgactgtgattgtgaatc | | |
| *Noc_F* | ggcaactcaagcgatgttca | Amplifies the *noc* locus of *S. aureus* | This study |
| *Noc_R* | acctttgaattgccagaaga | | This study |
| *lcpB_F* | tttcatttgataattgcctcaca | Amplifies the *lcpB* locus of *S. aureus* | This study |
| *lcpB_R* | ggagtgcttcatagttttctcg | | This study |
| *lcpC_F* | tagtaaaggagtggtgggat | Amplifies the *lcpC* locus of *S. aureus* | This study |
| *lcpC_R* | atcaccttctatttacgggc | | This study |
| *TnPbp3_F* | tgatgaaaacattacagtgaatg | Amplifies the *pbp3* locus of *S. aureus* | (Wacnik et al., 2022) |
| *TnPbp3_R* | gtatcgccatatggatattttc | | (Wacnik et al., 2022) |
| *ypsA_F* | acattgaacaactttctgcg | Amplifies the *ypsA* locus of *S. aureus* | This study |
| *ypsA_R* | actttgatcttcagaccact | | This study |
| *tarO_F* | gcttcgaacatgtctgaatcgactc | Amplifies the *tarO* locus of *S. aureus* | (Salamaga et al., 2021) |
| *tarO_R* | gcagttacctttcgatataccaactg | | (Salamaga et al., 2021) |
| *pbp4-1* | ctgcagaaaactttattttcaac | Amplifies a region of the *pbp4* locus of *S. aureus* | (Sutton et al., 2021) |
| *pbp4-5* | tatatagaactatcgatac taaac | | (Sutton et al., 2021) |
| pMAD_*divIVA_1* | cgttacacattaactagacaatcatcaatcgctgcaattatc | Amplifies 1000bp upstream of *divIVA* for insertion into pMAD by Gibson assembly | This Study |
| pMAD_*divIVA_2* | tatttaattcttgttatcctccttaaatcattac | | This Study |
| pMAD_*divIVA_3* | aggataacaagaattaaataaagacagacgc | Amplifies 1000bp downstream of *divIVA* for insertion into pMAD by Gibson assembly | This Study |
| pMAD_*divIVA_4* | gaattcgagctcccgggtacttttcgccatttacattg | | This Study |

| | | | |
|---|---|---|---|
| pKASBAR_*divIVA_F* | cagctatgaccatgattacgaagtgaatcacactattgttg | Primers to amplify *divIVA* and incorporate into pKASBAR via Gibson assembly. | This Study |
| pKASBAR_*divIVA_R* | ctgcccttttttttgccccggttacttcttagttgtttctgaatc | | This Study |
| pKASBAR_*gpsB_F* | ggaaacagctatgaccatgattacgtatttagtaattatttaccaaatacagct | Primers to amplify *gpsB* and incorporate into pKASBAR via Gibson assembly. | This Study |
| pKASBAR_*gpsB_R* | cgggatccggccatgtaggccagataggcaagtacaagtatgtgttgt | | This Study |
| piMAY_*gpsB_up_F* | agggaacaaaagctgggtactcaacaatagctttcttagttatcgcc | Primers to amplify 1000bp upstream of *gpsB* to incorporate into piMAY | This Study |
| piMAY_*gpsB_up_R* | tacgaggaattttactaaatacaaaagtttaactgtct | | This Study |
| piMAY_*gpsB_down_F* | ggttcgctggtttttccacctcattagaaactttga | Primers to amplify 1000bp downstream of *gpsB* to incorporate into piMAY | This Study |
| piMAY_*gpsB_down_R* | gctccaccgcggtggcggccttaaatcaaattatatagagtgtt | | This Study |
| pGL433_*kan_F* | gtatttagtaaaattcctcgtaggcgctcg | Primers to amplify the kanamycin resistance cassette from pGL433 to be incorporated into piMAY in between the *gpsB* upstream and downstream regions. | This Study |
| pGL433_*kan_R* | ggtggaaaaaccagcgaaccatttgaggtg | | This Study |
| pKB-*divIVA-F* | ctgcccttttttttgccccggatcatcaatcgctgcaattatc | Primers to amplify *divIVA-gfp* construct for incorporation into pKASBAR by Gibson assembly. | This Study |
| pKB-*divIVA-R* | cagctatgaccatgattacgttatttatacaattcgtccatacctaatg | | This Study |
| *Restriction sites are in capitals and overhangs for the Gibson assembly in blue | | | |

**Supplementary table 3 Oligonucleotides used in this study**

| | SH1000 | SH1000 *pbp3::ery* | SH1000 *gpsB::kan* | SH1000 *rodA::spec* | SH1000 *pbp3::ery gpsB::kan* | SH1000 *rodA::spec pbp3::ery* | SH1000 *rodA::spec gpsB::kan* | SH1000 *rodA::spec pbp3::ery gpsB::kan* |
|---|---|---|---|---|---|---|---|---|
| SH1000 | | * 0.0284 | **** <0.0001 | ns 0.1328 | **** <0.0001 | ** 0.0018 | **** <0.0001 | **** <0.0001 |
| SH1000 *pbp3::ery* | * 0.0284 | | *** 0.0010 | ns 0.9981 | * 0.0164 | ns 0.9971 | ns 0.0970 | *** 0.0002 |
| SH1000 *gpsB::kan* | **** <0.0001 | *** 0.0010 | | **** <0.0001 | ns 0.9930 | * 0.0102 | ns 0.8371 | ns >0.9999 |
| SH1000 *rodA::spec* | ns 0.1328 | ns 0.9981 | **** <0.0001 | | *** 0.0009 | ns 0.8649 | ** 0.0094 | **** <0.0001 |
| SH1000 *pbp3::ery gpsB::kan* | **** <0.0001 | * 0.0164 | ns 0.9930 | *** 0.0009 | | ns 0.1061 | ns 0.9983 | ns 0.9453 |
| SH1000 *rodA::spec pbp3::ery* | ** 0.0018 | ns 0.9971 | * 0.0102 | ns 0.8649 | ns 0.1061 | | ns 0.3864 | ** 0.0028 |
| SH1000 *rodA::spec gpsB::kan* | **** <0.0001 | ns 0.0970 | ns 0.8371 | ** 0.0094 | ns 0.9983 | ns 0.3864 | | ns 0.6249 |
| SH1000 *rodA::spec pbp3::ery gpsB::kan* | **** <0.0001 | *** 0.0002 | ns >0.9999 | **** <0.0001 | ns 0.9453 | ** 0.0028 | ns 0.6249 | |

**Supplementary Table 4 Statistics for Figure 3E**

Derived *p* values for Figure 3E analysed using a one-way ANOVA with multiple comparisons.

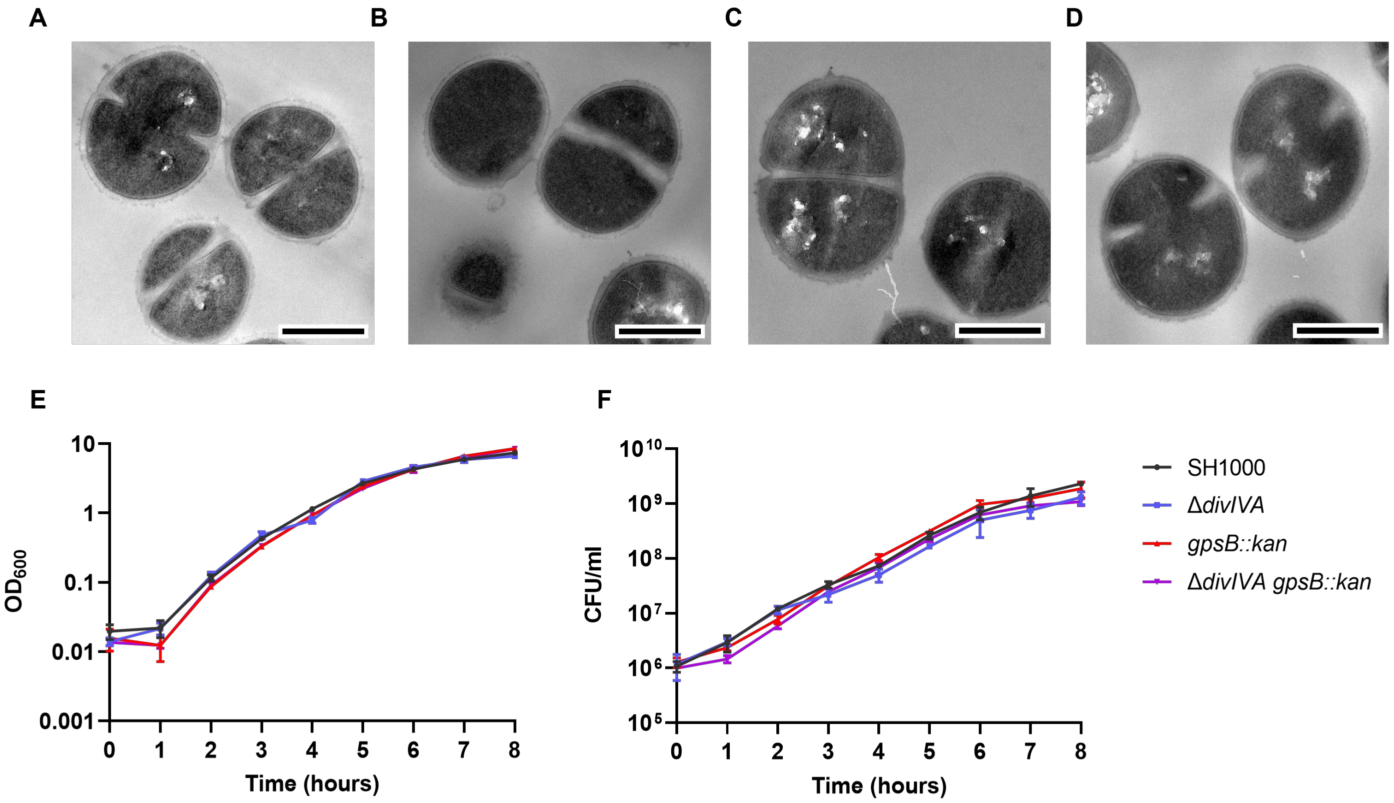


**Supplementary Figure 1** Morphology and growth analysis of *gpsB and divIVA* mutants

Representative thin section TEM micrographs for **(A)** SH1000, **(B)** SH1000 Δ*divIVA,* **(C)** SH1000 *gpsB::kan* and **(D)** SH1000 Δ*divIVA gpsB::kan* (scale bars represent 500 nm). **(E)** Growth curves and **(F)** viability of SH1000 (black lines), SH1000 Δ*divIVA* (blue lines)*,* SH1000 *gpsB::kan* (red lines) and SH1000 Δ*divIVA gpsB::kan* (purple lines). Bacterial cultures were prepared in triplicate and error bars show standard deviation.

**Video 1** The localisations of GpsB and DivIVA

Videos showing Z-stacks of widefield fluorescence microscopy and maximum intensity projections for SH1000 *gpsB::gpsB-mCherry kanR ΔdivIVA geh::divIVA-megfp*. GpsB-mCherry (magenta), DivIVA-GFP (yellow) and HADA (cyan) are shown. **(A)** and **(B)** show different representative examples.

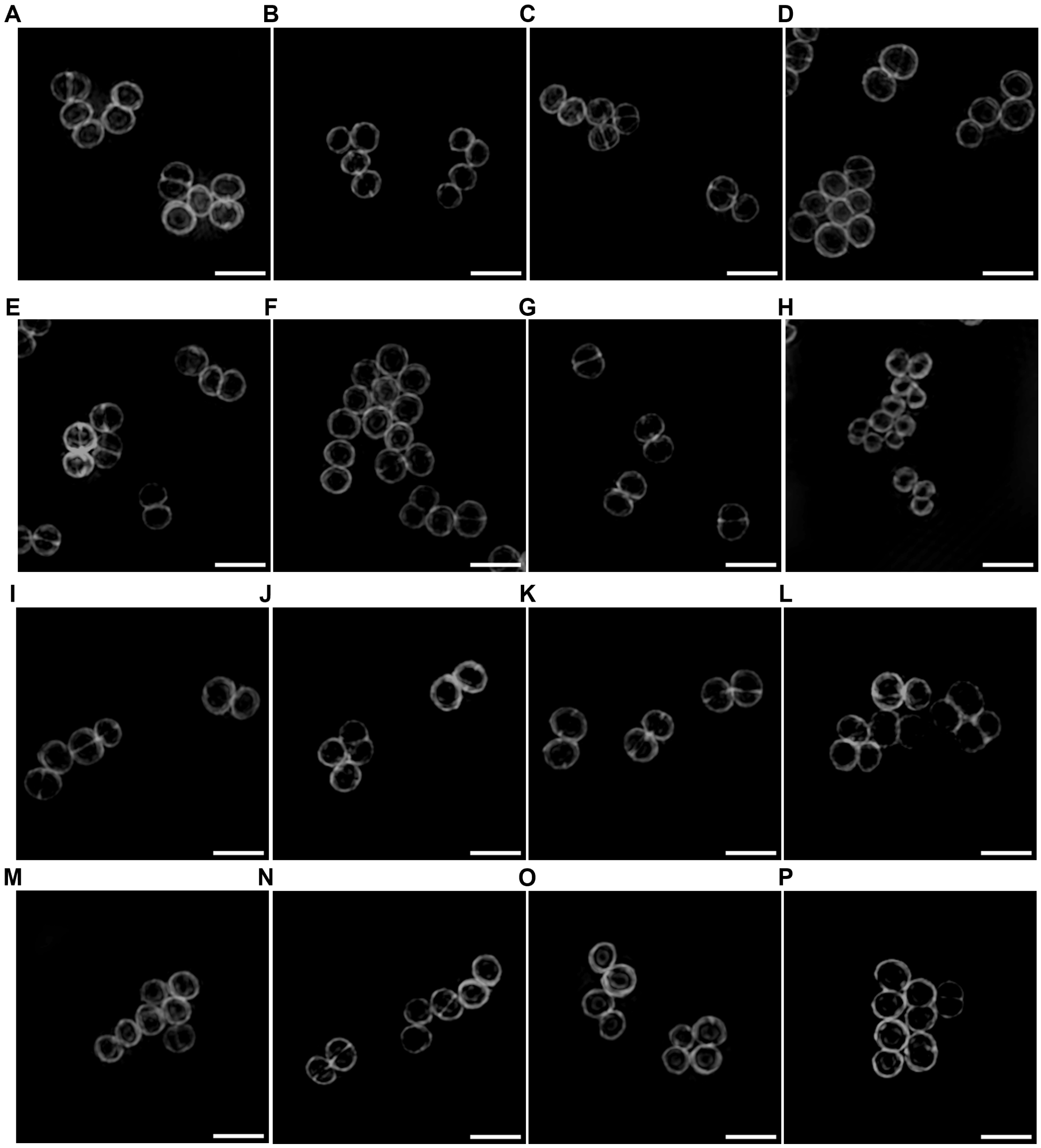


**Supplementary Figure 2** Representative SIM micrographs of strains

Representative images of SIM micrographs of *S. aureus* strains **(A)** SH1000 **(B)** SH1000 *ΔdivIVA gpsB::kan* **(C)** SH1000 *lcpC::ery* **(D)** SH1000 *ΔdivIVA gpsB::kan lcpC::ery* **(E)** SH1000 *pbp4::ery* **(F)** SH1000 *ΔdivIVA gpsB::kan pbp4::ery* **(G)** SH1000 *ypfP::ery* **(H)** SH1000 *ΔdivIVA gpsB::kan ypfP::ery* **(I)** SH1000 *ΔdivIVA* **(J)** SH1000 *ΔdivIVA lcpC::ery* **(K)** SH1000 *ΔdivIVA pbp4::ery* **(L)** SH1000 *ΔdivIVA ypfP::ery* **(M)** SH1000 *gpsB::kan* **(N)** SH1000 *gpsB::kan lcpC::ery* **(O)** SH1000 *gpsB::kan pbp4::ery* **(P)** SH1000 *gpsB::kan ypfP::ery* labelled with NHS Ester 555 (scale bars represent 2 μm).

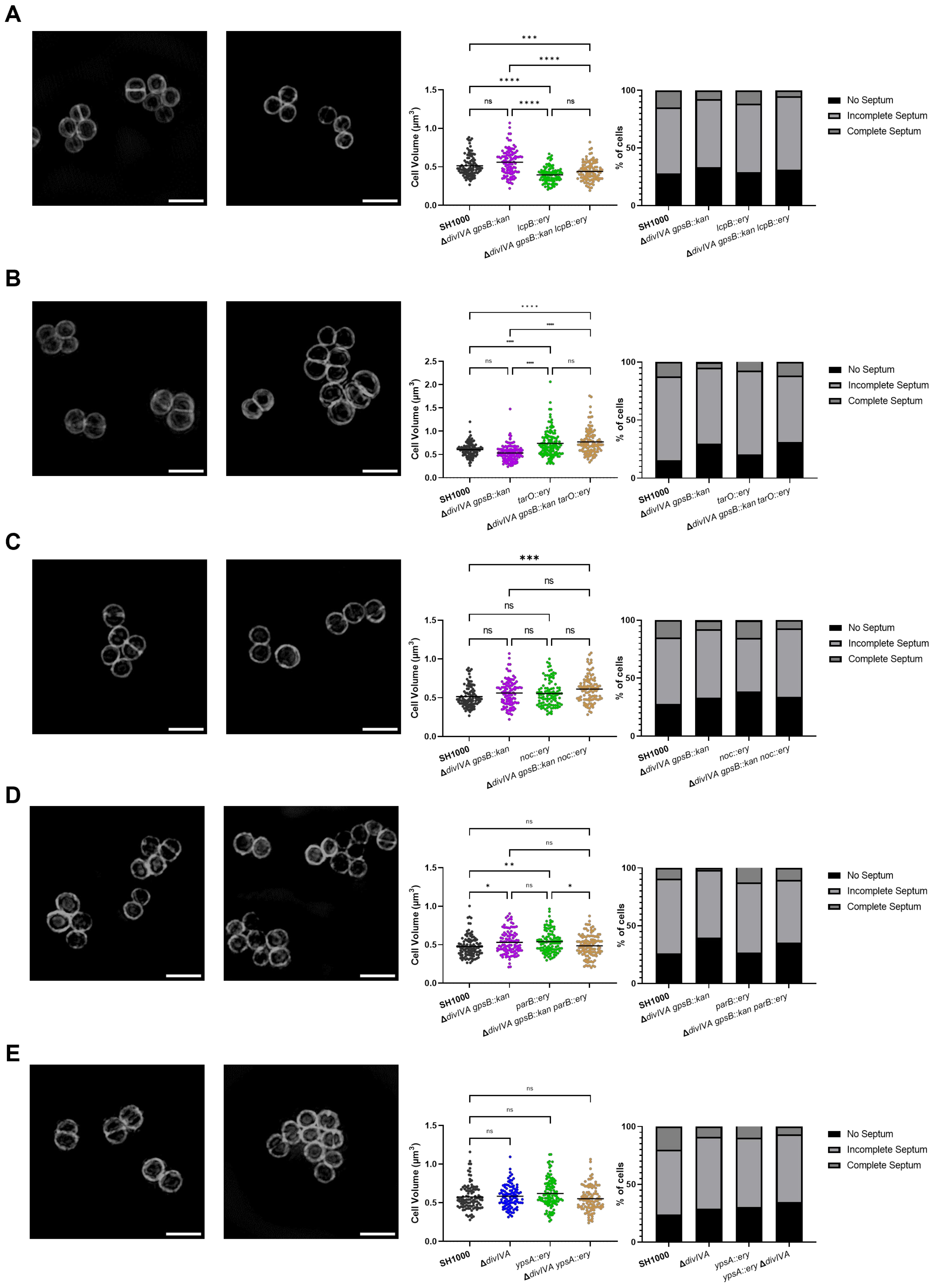


**Supplementary Figure 3** Phenotypes of *divIVA gpsB* combination mutants

SIM images of mutants of genes of interest and triple mutants of *divIVA* and *gpsB* with the target genes, with cell volume and the percentage of cells in specific stages of the cell cycle recorded from these compared to SH1000 and SH1000 *ΔdivIVA gpsB::kan*. **(A)** SH1000 *lcpB::ery* and SH1000

Δ*divIVA gpsB::kan lcpB::ery.* **(B)** SH1000 *tarO::ery* and SH1000 Δ*divIVA gpsB::kan tarO::ery.* **(C)** SH1000 *noc::ery* and SH1000 *noc::ery ΔdivIVA gpsB::kan.* **(D)** SH1000 *parB::ery* and SH1000 Δ*divIVA gpsB::kan parB::ery.* **(E)** SH1000 *ypsA::ery* and SH1000 Δ*divIVA ypsA::ery.* Results for cell volume analysis were analysed using a two-way ANOVA (* $p < 0.05$, ** $p < 0.005$, *** $p < 0.001$, **** $p < 0.0001$).

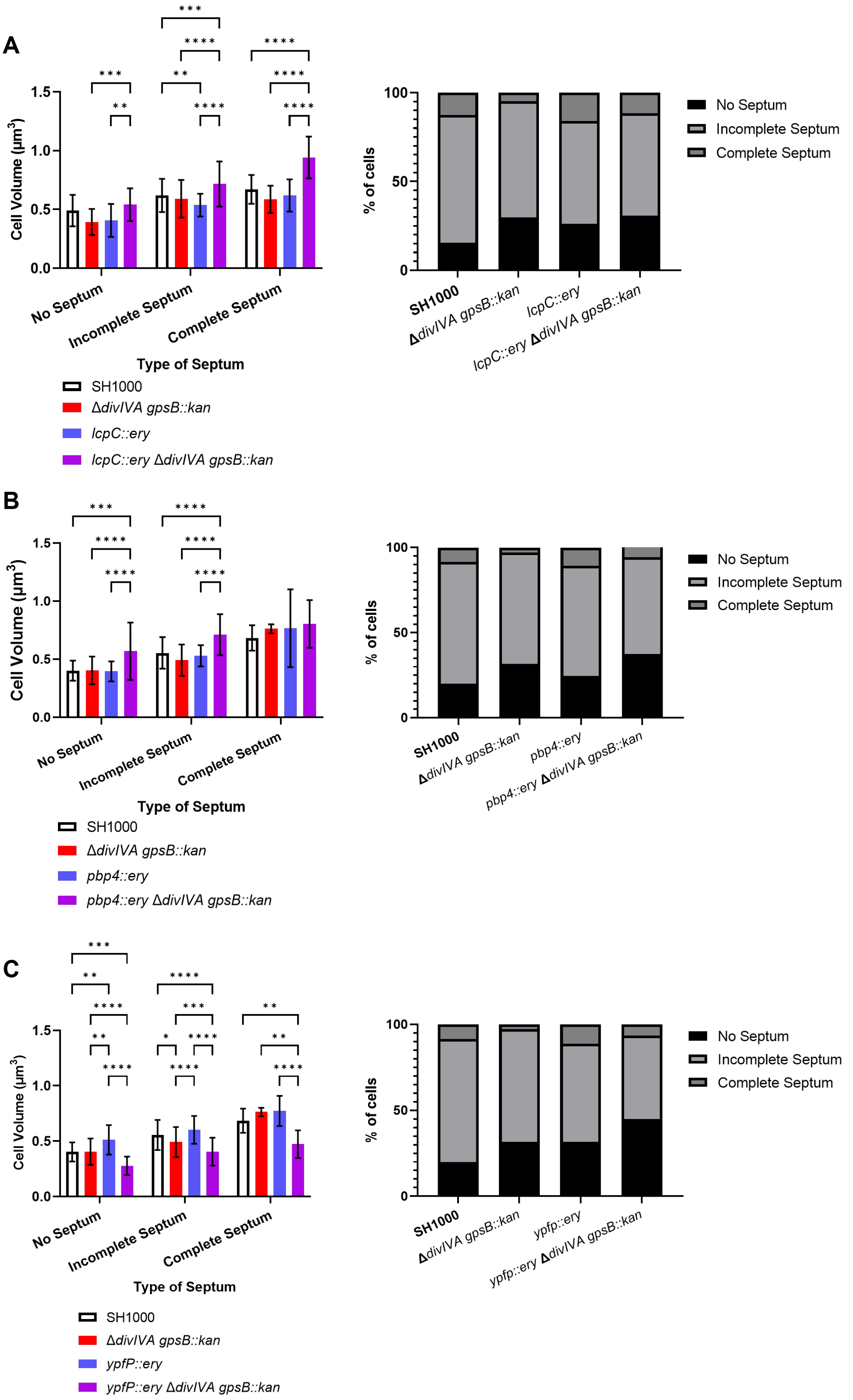


**Supplementary Figure 4** Phenotypic analysis of *divIVA gpsB* combination mutants

Cell volumes per stage of the cell cycle and percentage of cells in specific parts of the cell cycle based on septum completion. (A) SH1000 (black bars, n = 104), SH1000 *ΔdivIVA gpsB::kan* (red bars, n = 124, $p$ = 0.0028), SH1000 *lcpC::ery* (blue bars, n = 107), SH1000 *ΔdivIVA gpsB::kan lcpC::ery* (purple bars, n = 104). (B) SH1000 (black bars, n = 105), SH1000 *ΔdivIVA gpsB::kan* (red bars, n = 107, $p$ = 0.0028), SH1000 *pbp4::ery* (blue bars, n = 102), SH1000 *ΔdivIVA gpsB::kan pbp4::ery* (purple bars, n = 125). (C) SH1000 (black bars, n = 105), SH1000 *ΔdivIVA gpsB::kan* (red bars, n = 107), SH1000 *ypfP::ery* (blue bars, n = 113), SH1000 *ΔdivIVA gpsB::kan ypfP::ery* (purple bars, n = 117). Results were analysed using a two-way ANOVA (* $p < 0.05$, ** $p < 0.005$, *** $p < 0.001$, **** $p < 0.0001$).

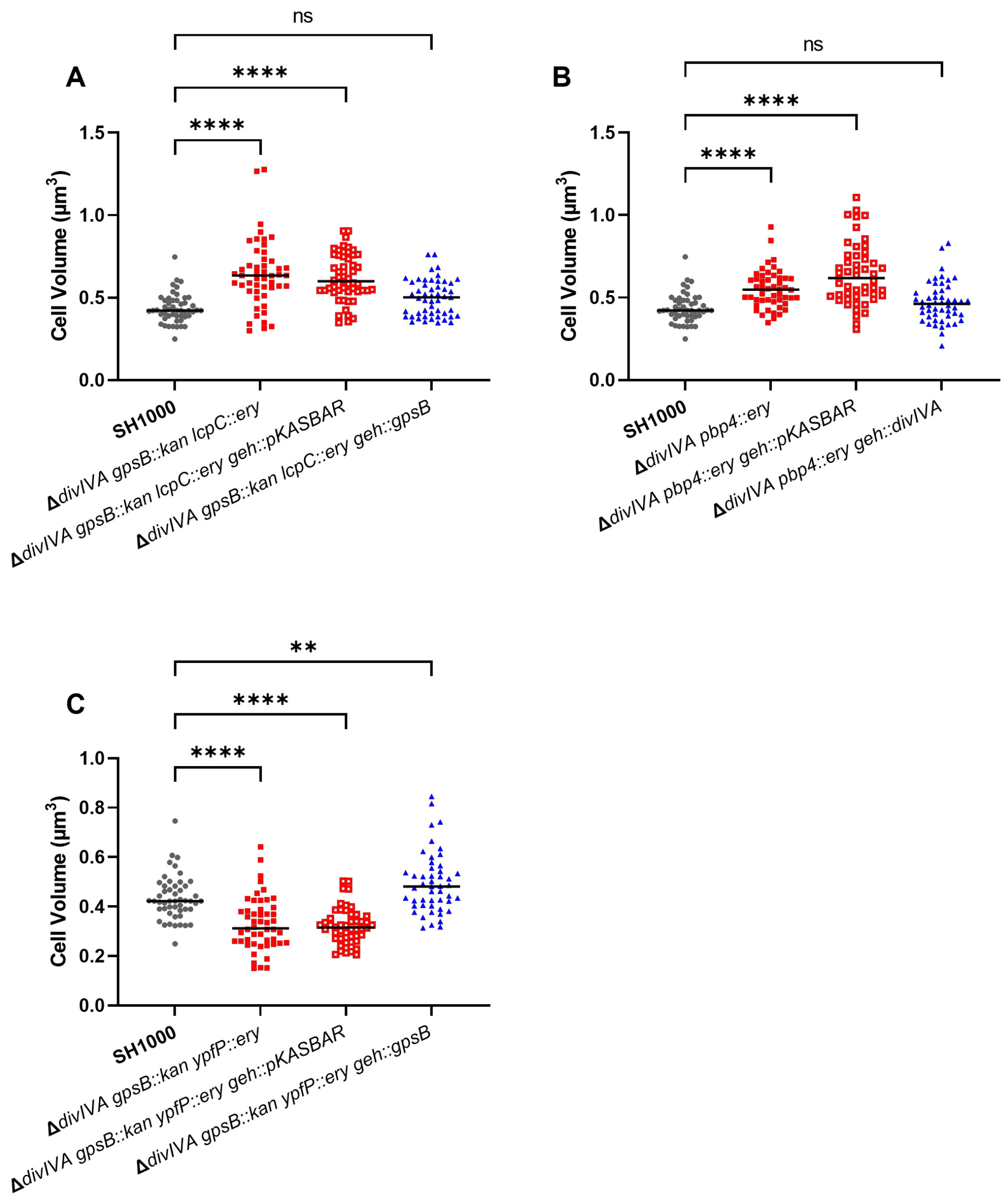


**Supplementary Figure 5** Complementation of strains of interest

Complementation and cell volume measurements for strains of interest from Figure 2. **(A)** SH1000 (Black circles, n = 50), SH1000 *ΔdivIVA gpsB::kan lcpC::ery* (red squares, n = 50, $p < 0.0001$), SH1000 *ΔdivIVA gpsB::kan lcpC::ery geh::pKASBAR* (open red squares, n = 53, $p < 0.0001$) and SH1000 *ΔdivIVA gpsB::kan lcpC::ery geh::gpsB* (blue triangles, n = 51, $p = 0.0638$). **(B)** SH1000 (Black circles, n = 50), SH1000 *ΔdivIVA pbp4::ery* (red squares, n = 50, $p < 0.0001$), SH1000 *ΔdivIVA pbp4::ery geh::pKASBAR* (open red squares, n = 50, $p < 0.0001$) and SH1000 *ΔdivIVA pbp4::ery geh::divIVA* (blue triangles, n = 50, $p = 0.4090$). **(C)** SH1000 (Black circles, n = 50), SH1000 *ΔdivIVA gpsB::kan ypfP::ery* (red squares, n = 50, $p < 0.0001$), SH1000 *ΔdivIVA gpsB::kan ypfP::ery geh::pKASBAR* (open red squares, n = 50, $p < 0.0001$) and SH1000 *ΔdivIVA gpsB::kan ypfP::ery geh::gpsB* (blue triangles, n = 49, $p = 0.0095$).

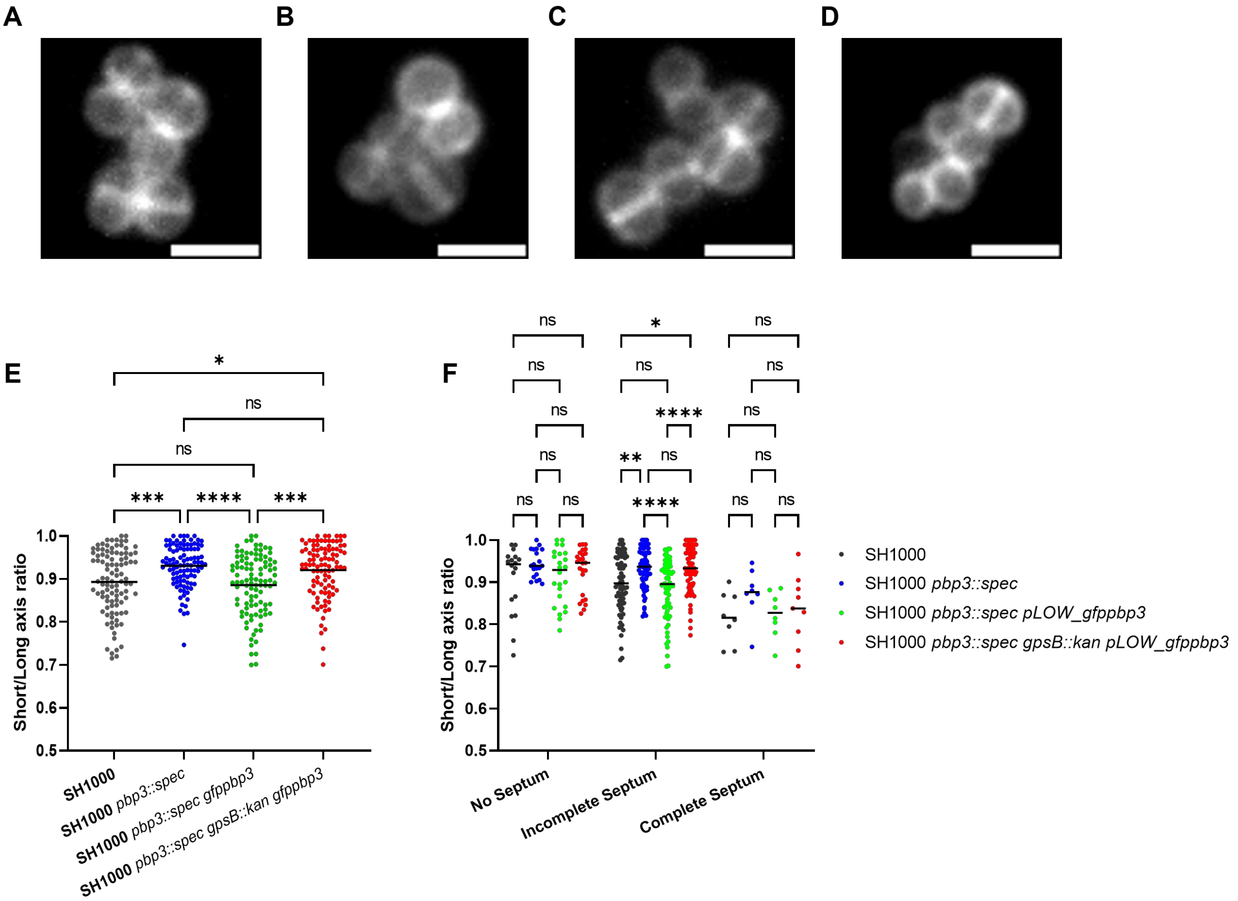


**Supplementary Figure 6 Functionality of the GFP-PBP3 fusion protein**

Micrographs for (A) SH1000 (B) SH1000 *pbp3::spec* (C) SH1000 *pbp3::spec gfppbp3* (D) SH1000 *pbp3::spec gfppbp3 gpsB::kan* each labelled with NHS Ester 555. Scale bars represent 2 µm. (E) The short/long axis ratio of SH1000 (black circles, n = 100), SH1000 *pbp3::spec* (blue circles, n = 101), SH1000 *pbp3::spec gfppbp3* (green circles, n = 103) and SH1000 *pbp3::spec gfppbp3 gpsB::kan* (red circles, n = 101). Data was analysed using a one-way ANOVA (* $p$ = 0.0134, ** $p$ = 0.0011, *** $p$ = 0.0003(SH1000 to SH1000 *pbp3::spec*) and 0.0007 (SH1000 *pbp3::spec gfppbp3* to SH1000 *pbp3::spec gpsB::kan gfppbp3*), **** < 0.0001). **(F)** The short/long axis ratios from **(E)** organised by the stage of the cell cycle cells were in ($p$ values * $p$ = 0.0110, ** $p$ = 0.0031, **** $p$ <0.0001). Analysed using a two-way ANOVA with multiple comparisons.

### References


Arnaud, M., Chastanet, A., and Débarbouillé, M. (2004). New vector for efficient allelic replacement in naturally nontransformable, low-GC-content, gram-positive bacteria. *Appl Environ Microbiol* 70, 6887–6891. doi: 10.1128/AEM.70.11.6887-6891.2004.

Boldock, E., Surewaard, B. G. J., Shamarina, D., Na, M., Fei, Y., Ali, A., et al. (2018). Human skin commensals augment Staphylococcus aureus pathogenesis. *Nat Microbiol* 3, 881–890. doi: 10.1038/s41564-018-0198-3.

Bottomley, A. L., Kabli, A. F., Hurd, A. F., Turner, R. D., Garcia-Lara, J., and Foster, S. J. (2014). Staphylococcus aureus DivIB is a peptidoglycan-binding protein that is required for a morphological checkpoint in cell division. *Molecular Microbiology* 94, 1041–1064. doi: 10.1111/mmi.12813.

Fey, P. D., Endres, J. L., Yajjala, V. K., Widhelm, T. J., Boissy, R. J., Bose, J. L., et al. (2013). A genetic resource for rapid and comprehensive phenotype screening of nonessential Staphylococcus aureus genes. *mBio* 4, e00537-00512. doi: 10.1128/mBio.00537-12.

Horsburgh, M. J., Aish, J. L., White, I. J., Shaw, L., Lithgow, J. K., and Foster, S. J. (2002). sigmaB modulates virulence determinant expression and stress resistance: characterization of a functional rsbU strain derived from Staphylococcus aureus 8325-4. *J Bacteriol* 184, 5457–5467. doi: 10.1128/JB.184.19.5457-5467.2002.

Kreiswirth, B. N., Löfdahl, S., Betley, M. J., O'Reilly, M., Schlievert, P. M., Bergdoll, M. S., et al. (1983). The toxic shock syndrome exotoxin structural gene is not detectably transmitted by a prophage. *Nature* 305, 709–712. doi: 10.1038/305709a0.

Liew, A. T. F., Theis, T., Jensen, S. O., Garcia-Lara, J., Foster, S. J., Firth, N., et al. (2011). A simple plasmid-based system that allows rapid generation of tightly controlled gene expression in Staphylococcus aureus. *Microbiology* 157, 666–676. doi: 10.1099/mic.0.045146-0.

Monk, I. R., Shah, I. M., Xu, M., Tan, M.-W., and Foster, T. J. (2012). Transforming the untransformable: application of direct transformation to manipulate genetically Staphylococcus aureus and Staphylococcus epidermidis. *mBio* 3, e00277-11. doi: 10.1128/mBio.00277-11.

Salamaga, B., Kong, L., Pasquina-Lemonche, L., Lafage, L., von Und Zur Muhlen, M., Gibson, J. F., et al. (2021). Demonstration of the role of cell wall homeostasis in Staphylococcus aureus growth and the action of bactericidal antibiotics. *Proc Natl Acad Sci U S A* 118, e2106022118. doi: 10.1073/pnas.2106022118.

Sutton, J. A. F., Carnell, O. T., Lafage, L., Gray, J., Biboy, J., Gibson, J. F., et al. (2021). Staphylococcus aureus cell wall structure and dynamics during host-pathogen interaction. *PLOS Pathogens* 17, e1009468. doi: 10.1371/journal.ppat.1009468.

Tinajero-Trejo, M., Carnell, O., Kabli, A. F., Pasquina-Lemonche, L., Lafage, L., Han, A., et al. (2022). The Staphylococcus aureus cell division protein, DivIC, interacts with the cell wall and controls its biosynthesis. *Commun Biol* 5, 1–13. doi: 10.1038/s42003-022-04161-7.

Wacnik, K., Rao, V. A., Chen, X., Lafage, L., Pazos, M., Booth, S., et al. (2022). Penicillin-Binding Protein 1 (PBP1) of Staphylococcus aureus Has Multiple Essential Functions in Cell Division. *mBio* 0, e00669-22. doi: 10.1128/mbio.00669-22.

Wheeler, R., Turner, R. D., Bailey, R. G., Salamaga, B., Mesnage, S., Mohamad, S. A. S., et al. (2015). Bacterial Cell Enlargement Requires Control of Cell Wall Stiffness Mediated by Peptidoglycan Hydrolases. *mBio* 6, e00660. doi: 10.1128/mBio.00660-15.
